# Rac1, Rac3 GTPases and TPC2 are required for axonal outgrowth and migration of cortical interneurons

**DOI:** 10.1101/2022.11.07.515393

**Authors:** Zouzana Kounoupa, Simona Tivodar, Kostas Theodorakis, Dimitrios Kyriakis, Myrto Denaxa, Domna Karagogeos

**Author notes:** Address correspondence to Prof. Domna Karagogeos, IMBB, PO Box 1385, Vassilika Vouton, Heraklion, 71110 Crete, Greece.

## Abstract

Rho GTPases, among them Rac1 and Rac3, are major transducers of extracellular signals and are involved in multiple cellular processes. In cortical interneurons, the neurons that control excitation/inhibition balance of cortical circuits, Rac1 and Rac3 are essential for their development. Ablation of both, leads to a severe reduction in the numbers of mature interneurons found in the murine cortex, which is partially due to abnormal cell cycle progression of interneuron precursors and defective formation of their growth cones. Here we present new evidence that upon Rac1 and Rac3 ablation, centrosome, Golgi complex and lysosome positioning are significantly perturbed, thus affecting both interneuron migration and axon growth. Moreover, for the first time we provide evidence of altered expression and localization of the two-pore channel 2 (TPC2) voltage-gated ion channel that mediates Ca^2+^ release. Pharmacological inhibition of TPC2 negatively affected axonal growth and migration of interneurons. Our data taken together suggest that TPC2 contributes to the severe phenotype in axon growth initiation, extension and interneuron migration in the absence of Rac1 and Rac3.

**SUMMARY STATEMENT:** Rac1/3 severely affect cortical interneuron migration by affecting centrosome, Golgi and lysosome positioning. TPC2 likely contributes to the phenotype by decreasing axonogenesis and somatic migration.

## Introduction

Cortical GABAergic interneurons (CINs) play important roles in cortical function, from fine-tuning the excitability of pyramidal neurons to controlling the excitation/inhibition balance. Their loss or dysfunction has been implicated in severe disorders such as schizophrenia, epilepsy, autism spectrum disorders and others (Marín, 2012). CINs are a highly diverse neuronal population deriving from well-defined domains of the ventral telencephalon. The majority of them originates from the medial ganglionic eminence (MGE) and migrate tangentially, following specific migratory routes to populate the developing cortex (Lim et al., 2018; Marín and Rubenstein, 2001; Marín and Rubenstein, 2003). During their tangential migration, interneurons follow a stereotypic cyclic type of movement (Bellion et al., 2005). Efforts to uncover the intracellular mediators responsible for the response of CINs to environmental cues, have demonstrated that Rho-GTPases play significant roles in interneuron development through diverse functions (Corbetta et al., 2005; Corbetta et al., 2009; de Curtis, 2014; Kalemaki et al., 2018; Kounoupa and Karagogeos, 2022; Pennucci et al., 2016; Tivodar et al., 2015; Vidaki et al., 2012).

Rho-GTPases are integrators of multiple extracellular signals and are required for many biological functions in many cell types, including cytoskeleton organization, cell morphogenesis, transcription, cell cycle progression and apoptosis (Jaffe and Hall, 2005; Scala et al., 2021). Moreover, Rho-GTPases are shown to act in axon pathfinding and migration processes from worms and flies to mammalian neurons (Gonzalez-Billault et al., 2012; Hakeda-Suzuki et al., 2002; Lundquist, 2003; Lundquist et al., 2001). Rho GTPases affect both the actin cytoskeleton as well as the organization of microtubules (Etienne-Manneville and Hall, 2001; Palazzo et al., 2001; Van Aelst and D’Souza-Schorey, 1997). Both of these cytoskeletal components are crucial in various aspects of neuronal development, including migration, axon formation and outgrowth, as well as dendrite and dendritic spine formation and maintenance (Coles and Bradke, 2015).

Previous work from our lab showed that mice lacking Rac1 exclusively in MGE-derived cells, exhibit a 50% reduction in the number of GABAergic interneurons found in the mature cortex. Mutant CINs aggregated in the ventral telencephalon and displayed affected growth cones and abnormal axon extension (Vidaki et al., 2012), leading to the conclusion that Rac1 is required cell autonomously for CIN development and cell cycle progression. We have subsequently shown that double ablation of Rac1/Rac3, leads to a failure for axon extension beyond a certain length, defective growth cones and to an even more severe absence of CINs in the postnatal cortex up to 80% (Tivodar et al., 2015). In addition, proteins of the PAK-interacting exchange factor (PIX)/G-protein coupled receptor kinase-interacting protein (GIT) network expressed in MGE-derived cells and functionally linked to Rac1/3 have been shown to be involved in proper neuritic development (Franchi et al., 2016).

Here we provide evidence on the migratory behavior of CINs upon double ablation of Rac1/Rac3. Key steps and elements of the proper progression of the migratory cycle, such as centrosome and Golgi complex subcellular localization, are defective. Moreover, initiation of axon outgrowth and axonal transport, as evidenced by lysosomal locomotion patterns, are significantly impacted. RNA sequencing revealed genes with altered expression between wild type and double mutant CINs, among which, the two-pore channel 2 (TPCN2 encoding TPC2), a member of the superfamily of voltage-gated ion channels, that is mainly located in late endosomes/lysosomes and mediates Ca^2+^ release (Marchant and Patel, 2015; Patel and Kilpatrick, 2018). In Rac1/2 double mutant interneurons TPC2 protein levels are reduced and its subcellular localization is affected. Finally, pharmacological inhibition of TPC2 negatively affects axonal growth and migration of MGE-derived interneurons, similarly to the ablation of Rac1/3. We therefore propose that TPC2 is a novel Rac1/3 potential mediator of CIN development.

## Results

### Rac1 and Rac3 ablation affects the migratory behavior of the MGE-derived interneurons

We have previously shown that Rac1/Rac3dmut mice exhibit a dramatic loss of MGE derived interneurons in the developing cortex resulting from a cell autonomous delay in the cell cycle exit and defective migration (Tivodar et al., 2015). In order to elucidate the mechanisms underlying the described migratory defects, we performed real time imaging, for ∼8hrs, on acute brain slices from Rac1/Rac3wt and Rac1/Rac3dmut embryos, and recorded the migratory behavior of interneurons at the level of the corticostriatal boundary (Fig. 1A).

**Fig. 1.**
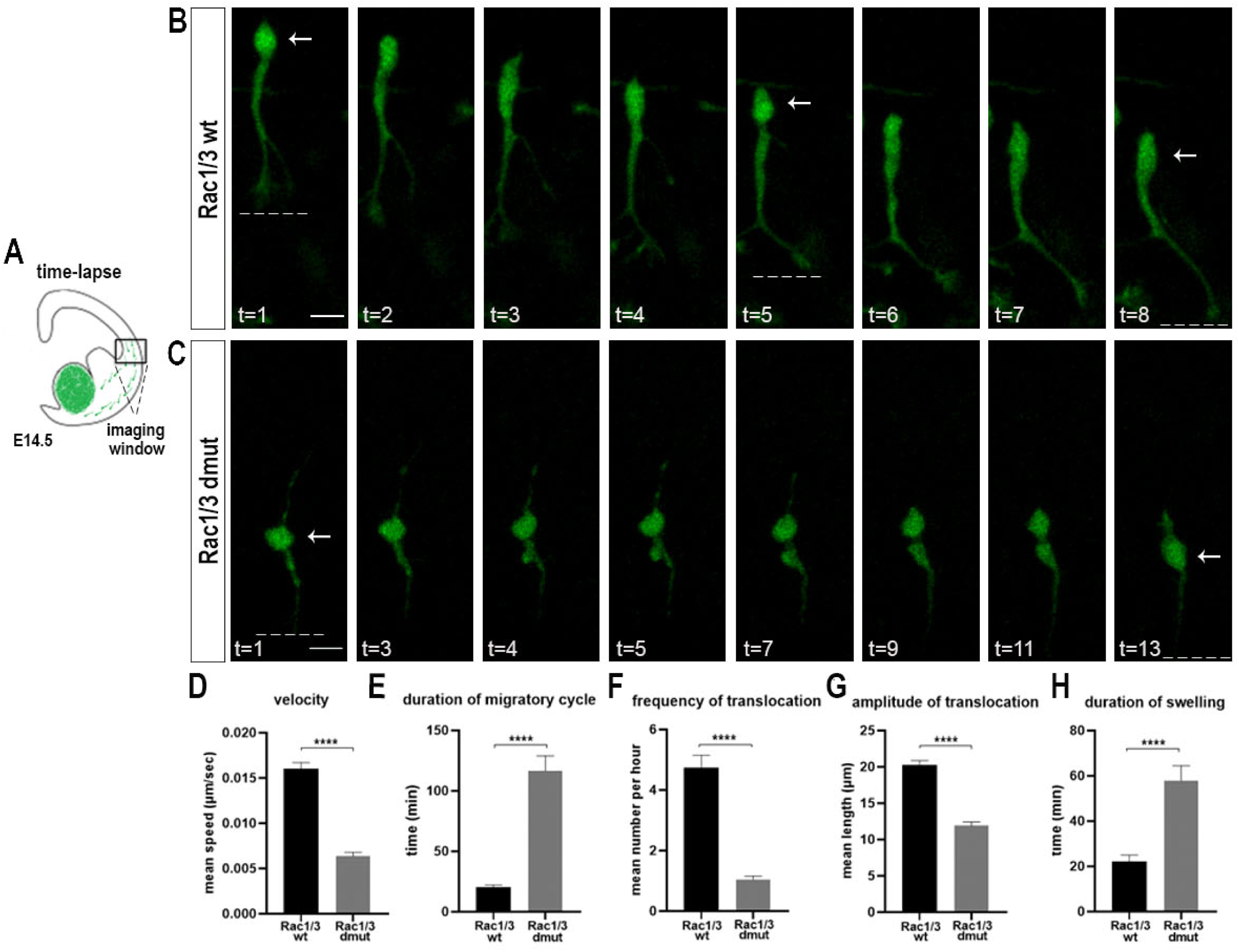
Ablation of Rac1 and Rac3 affects the migratory behavior of the MGE-derived interneurons. (A) Schematic representation of an E14.5 brain slice prepared for real-time recording of interneuron migration (box indicating the imaging window) (B) Time-lapse sequence of a migrating YFP-expressing Rac1/3wt interneuron (C) and a Rac1/3dmut interneuron. White arrows and dotted lines illustrate the beginning and ending position of the nucleus and the tip of the leading process respectively, during a migration cycle. Time-lapse images acquired every 3min for ∼8hrs. Histograms of (D) the migration velocity (mean±SEM: 0.01603±0.00069 for Rac1/3wt, 0.006359±0.00043 for Rac1/3dmut), (n=39-68 cells per condition, from 3-5 independent experiments) (E) the duration of the migratory cycle (mean±SEM: 20.59±1.53 Rac1/3wt, 116.6±12.37 Rac1/3dmut), (n=35-52 cells per condition, from 3-5 independent experiments), (F) the frequency of nuclear translocation of migrating interneurons (mean±SEM: 20.22±0.65 Rac1/3wt, 11.89±0.51 Rac1/3dmut), (n=19-25 cells per condition, from 3-5 independent experiments), (G) the amplitude of nuclear translocation (mean±SEM: 4.733±0.41 Rac1/3wt, 1.038±0.12 Rac1/3dmut), (n=21-28 cells per condition, from 3-5 independent experiments), (H) duration of swelling presence (n=34-49 cells per condition, from 3-5 independent experiments). Student’s t test, ****P < 0.0001. Error bars indicate SEM. Scale bars represent 10 μm.

During their migratory phase, interneurons exhibit a cyclic, three-step movement. At the beginning, a branched leading process that will become the axon, extends, followed by the formation of a swelling where the centrosome and Golgi apparatus translocate. Next the nucleus moves forward, into the swelling, in a saltatory manner and finally, the trailing process is retracted from the cell rear (Bellion, 2005). We observed the swelling formation protruding in front of the nucleus in the migratory cycle, followed by the saltatory nuclear translocation in both genotypes, even though only few Rac1/Rac3dmut interneurons entered the dorsal telencephalon (movie 1). However, we noticed that several other aspects of their behavior were affected (Fig. 1B, C). There was an approximately 2.5fold reduction in the mean velocity of the migrating interneurons in the Rac1/Rac3dmut (Fig. 1D) and the duration of the migratory cycle was significantly increased (Fig. 1E). Concomitantly, the frequency of nuclear translocation was significantly reduced (Fig. 1F), as was the mean amplitude of translocation in the Rac1/Rac3dmut interneurons (Fig. 1G). Lastly, we observed that the duration of the appearance of the swelling formation per migratory cycle was significantly increased in the absence of Rac1/Rac3 (Fig. 1H).

We then performed focal electroporation in embryonic brain slices from control and mutant embryos with an RFP vector, in order to label individual interneurons and follow, with time lapse imaging, their migratory behavior within the MGE, 24hrs after electroporation (Fig. S1A). In accordance with our previous observations (Fig. 1) Rac1/Rac3dmut interneurons exhibited significantly reduced motility. In some cases, we could detect the swelling formation but it was not followed by a nuclear translocation event during the monitoring time (Fig. S1B).

Collectively, these data demonstrate that the migratory dynamics are severely impaired due to the ablation of Rac1/Rac3 in CINs.

### Centrosome and Golgi complex positioning, as well as nucleokinesis, are affected upon ablation of Rac1/ Rac3 GTPases

Given the above observations, we wanted to address the positioning of the centrosome and the Golgi complex in Rac1/Rac3dmut interneurons, since previous studies have implicated both organelles in the regulation of neuronal migration (Koizumi et al., 2006; Tanaka et al., 2004; Tsai et al., 2007; Yanagida et al., 2012). For this, we cultured CINs on collagen-coated coverslips and after 3DIV we immunolabeled the centrosome and the Golgi complex for pericentrin and golgin 97, respectively (Fig. 2A-B’, E-F’). We quantified the distance of the centrosome from the nucleus and found that it was significantly reduced in the Rac1/Rac3dmut compared to control interneurons (Fig. 2C). Next, we categorized the cells according to this distance, in four distinct groups: cells with their centrosome present 0-2μm from the nucleus, 2-4μm, over 4μm and a group termed as “nucleus” where in a two-dimensional image, centrosome and DAPI staining were overlapping. As shown in Fig. 2D, Rac1/Rac3dmut CINs tend to localize their centrosome closer to the nucleus, compared to control cells. Similarly, the distance of the Golgi complex in these interneurons was significantly reduced (Fig. 2G). Using the same categorization and comparison with control cells, we observed that in about twice as many of the Rac1/Rac3dmut interneurons, the Golgi complex was located in close proximity to the nucleus and fewer dmut cells had their Golgi complex positioned in a distance higher than 4μm (Fig. 2H). Together these results revealed that the positioning of the centrosome and the Golgi complex were significantly affected in CINs upon ablation of Rac1/Rac3. Both were located closer to the nucleus *in vitro,* providing a correlation with the displayed reduction in the amplitude of translocation during nucleokinesis.

**Fig. 2.**
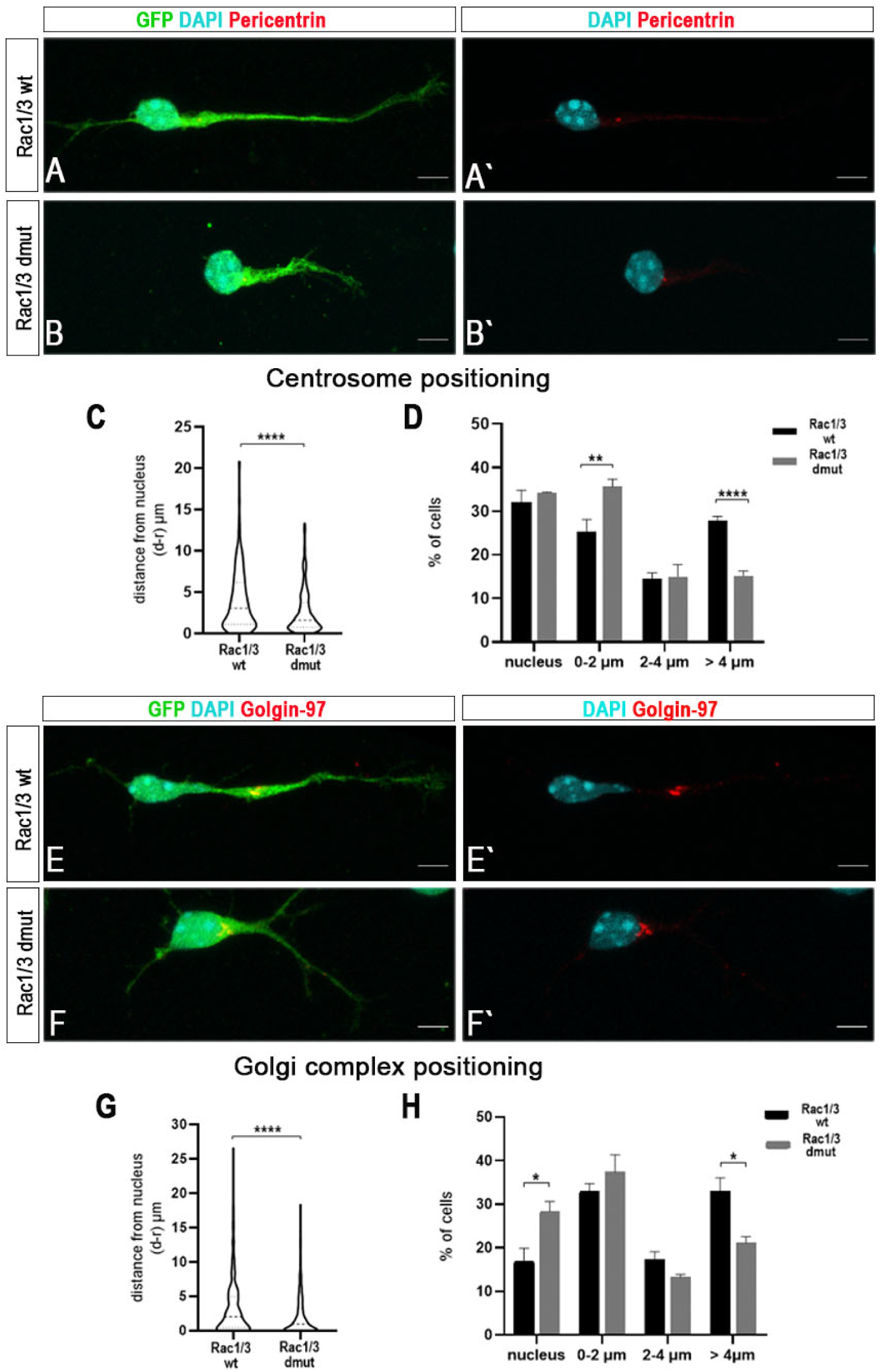
Centrosome and Golgi Complex positioning is altered due to the absence of Rac1 and Rac3 proteins in MGE-derived interneurons. (A-B’) Immunostaining of the Centrosome (pericentrin in red) and YFP (green) in cultured MGE-derived interneurons from E14.5 Rac1/3wt (A,À) and Rac1/3dmut (B,B’) embryos, 3DIV. The nucleus is labeled with DAPI (cyan). (C) Graph indicating the distance of the centrosome from the nucleus. (D) Histogram showing the percentage of cells with overlapping pericentrin-DAPI staining (noted as “nucleus”), or with centrosome located 0-2μm, 2-4μm and in distance higher than 4μm, from the nucleus. (E-F’) Immunostaining of Golgi complex (golgin-97 in red) and YFP (green) in cultured MGE-derived interneurons from E14.5 Rac1/3wt (E,È) and Rac1/3dmut (F,F’) embryos, 3DIV. The nucleus is labeled with DAPI (cyan). (G) Graph indicating the distance of the Golgi complex from the nucleus. (H) Histogram showing the percentage of cells with overlapping golgin-97-DAPI staining (noted as “nucleus”) or with Golgi complex located 0-2μm and 2-4μm or in distance higher than 4μm from the nucleus (n=90-120 cells per condition, from 3 independent experiments). Multiple t test, *P < 0.05, **P<0.01, ****P < 0.0001. Error bars indicate SEM. Scale bars represent 5 μm.

Optimal microtubule dynamics are necessary for the translocation of the nucleus (Garcez et al., 2015; Marín et al., 2010). Hence, we cultured MGE explants from Rac1/Rac3dmut and control mouse embryos, and labeled the microtubule cage that surrounds the nucleus for tyrosinated tubulin (ty-tubulin) and also the stable microtubules for acetylated tubulin (ac-tubulin), in migrating interneurons, 48h after plating (Fig. S1C). By measuring fluorescent intensity in the nuclear region, we found that there was a significant reduction of ac-tubulin and an increase in ty-tubulin in Rac1/Rac3dmut interneurons. The ratio between the two types of tubulin indicates a difference in microtubule dynamics upon ablation of Rac1/Rac3 that might negatively affect nucleokinesis (Fig. S1D-F).

Apart from microtubule dynamics and proper centrosome positioning, nuclear movement in migrating interneurons is additionally determined by the actomyosin dynamics, providing pushing forces at the rear of the cell (Bellion et al., 2005; Schaar and McConnell, 2005a). Actomyosin contractility in the leading process further contributes to the traction force required for the forward translocation and is regulated by RhoA signaling (Shinohara et al., 2012; Solecki et al., 2009). We asked whether this set of forces might also be affected in interneurons due to the loss of Rac1/Rac3. To address this question we stained migrating interneurons in brain slices from control and Rac1/Rac3dmut embryos, for phospho-Myosin Light Chain (pMLC). We noticed an accumulation of the pMLC in the perinuclear region in the Rac1/Rac3wt interneurons, while the pMLC signal was reduced in the Rac1/Rac3dmut cells (Fig. S2A). To further examine the defect, we cultured MGE-derived interneurons from Rac1/Rac3wt and Rac1/Rac3dmut embryos and we immunolabeled the cells using antibodies against pMLC, RhoA and pRhoA (Fig. S2B, E). Quantification of the pMLC fluorescent intensity revealed a significant reduction in the Rac1/Rac3dmut interneurons (Fig. S2C), especially in the region close to nucleus and in the axon, 5.96 μm from the nucleus up to its tip. In addition, we could observe an almost homogenous reduction of the signal throughout the Rac1/Rac3dmut cells (Fig. S2D). We also observed a significant reduction in the RhoA and pRhoA fluorescent intensity in the Rac1/Rac3dmut interneurons (Fig. S2G, H). Interestingly, the activation state of the protein was altered, as in the Rac1/Rac3dmut interneurons the pRhoA/RhoA ratio was significantly reduced in the region close to nucleus (Fig. S2F). Finally, we wanted to test whether a spatial difference in the activation state of the RhoA could be observed within the cells. Therefore, we quantified the pRhoA/RhoA ratio in standard intervals starting from the nucleus towards the principal neurite tip. A significant reduction was detected only in the regions close to the nucleus (Fig. S2I). Together these data suggest that in the absence of the two GTPases, the forces pushing the nucleus forward are affected. Moreover, western blot analysis performed on MGEs dissected from wt and Rac1mut embryos revealed increased F-actin in the latter genotype, indicating an alteration in actin cytoskeleton turnover (Fig. S2J-K).

In conclusion, ablation of Rac1 and Rac3 in CINs, results in significant defects in centrosome and Golgi complex subcellular localization and impairments in actomyosin contractility, leading to defective migratory behavior.

### Axon outgrowth is severely impaired in interneurons lacking Rac1 and Rac3

Migrating interneurons exhibit a bipolar morphology with a highly dynamic leading process (axon) that branches during the migratory cycle and a short trailing process (Arimura and Kaibuchi, 2007; Marín and Rubenstein, 2003; Marín et al., 2010). Formation and extension of the axon is a fundamental step for the migration thus we were interested in identifying how the ablation of Rac1/Rac3 might affect the axon outgrowth of CINs. We have previously described that the growth cones in double mutant interneurons exhibit extensive splitting of the leading process, absence of a formed axonal growth cone and impairments in lamellipodia formation (Tivodar et al., 2015). Immunostaining for Tuj-1 and F-actin in 2DIV cultured interneurons revealed additionally that filopodia were invaded by microtubules in the Rac1/Rac3dmut growth cones (Fig 3 A-F΄). To monitor axon outgrowth, we cultured dissociated cells from MGEs of both genotypes on Matrigel (Fig. 3G). We observed by live imaging that the rate of outgrowth in the Rac1/Rac3dmut was reduced. A more pronounced difference was observed in the axon extension rate between 2-4 hrs (Fig 3 H). By following the axon length for a period of 6 days (on days 2, 4 and 6) we discovered that the reduced growing rate was sustained in the Rac1/Rac3dmut interneurons compare to control ones, and the maximum length that the axons reached was significantly reduced (Fig 3 I). Sholl analysis at 6DIV revealed an over-branched phenotype, while the total length of the neurites was significantly decreased in Rac1/Rac3dmut interneurons (Fig 3 L, M). Hence, upon ablation of Rac1/Rac3 the morphology of CINs growth cones is severely disturbed resulting in over-branched neurons with defective axon initiation and axon elongation processes.

**Fig. 3.**
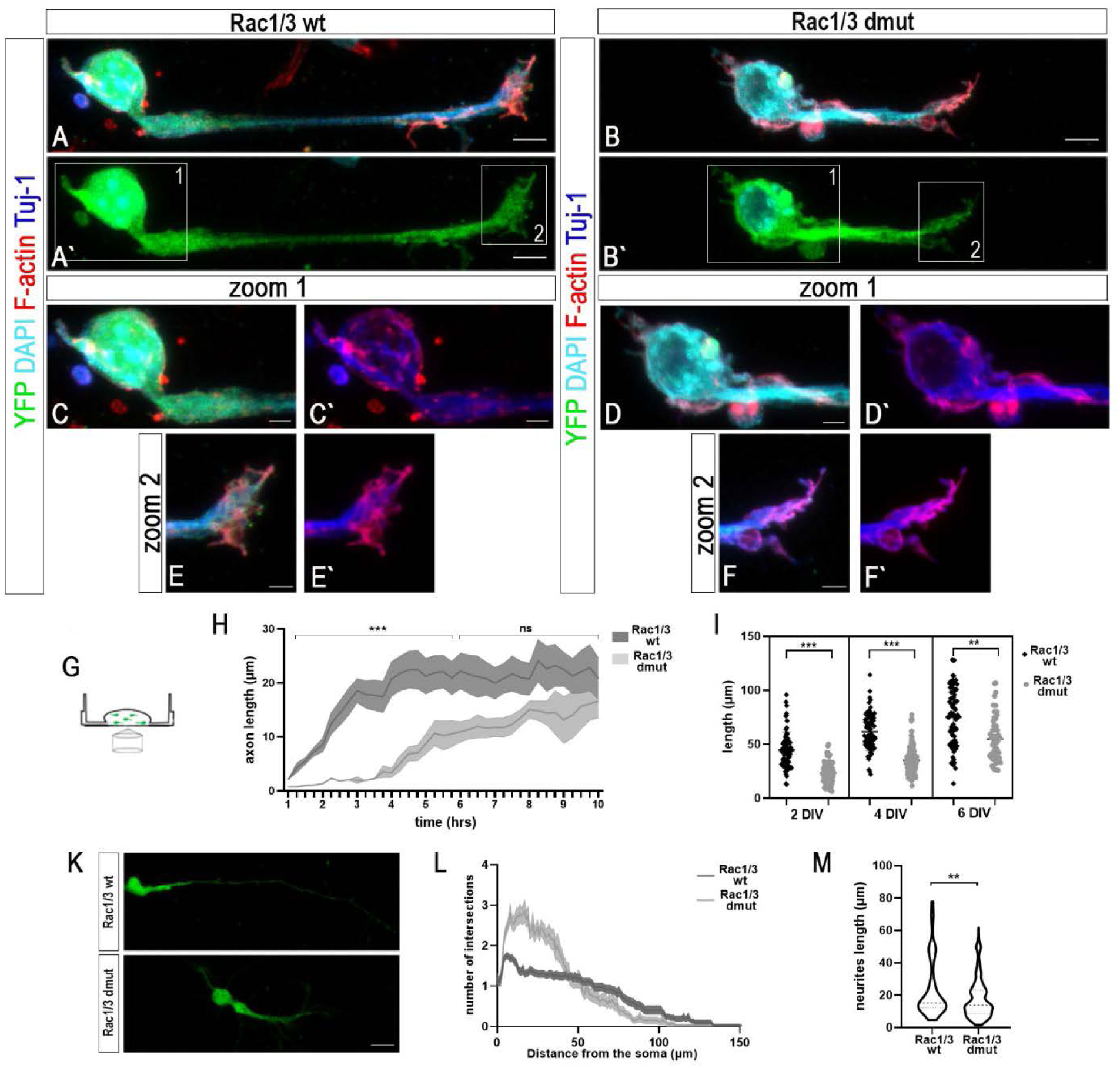
Ablation of Rac1 and Rac3 affects the axon growth of MGE-derived interneurons. Immunostaining for microtubules (Tuj-1, in blue), F-actin (phalloidin, in red), YFP (green) in cultured MGE-derived interneurons from E14.5 Rac1/3wt (A,À) and Rac1/3dmut (B,B’) embryos, 2DIV. The nucleus is labeled with DAPI (cyan) (C-D’). High magnification view of the boxed area 1, in the cell body of Rac1/3wt (C,C΄) and Rac1/3dmut interneuron (D,D’) and of the boxed area 2 (Ε-F’), in the growth cone of Rac1/3wt (E-È) and Rac1/3dmut interneuron (F-F’). (G) Time-lapse imaging of MGE-derived interneurons cultured on matrigel. (H) Graph indicating principal neurite outgrowth rate in the first 8hrs after plating of Rac1/3wt and Rac1/3dmut interneurons (I) Graph showing the length of the principle neurite 2,4,6 days after plating. (K) Immunostaining for YFP (green) of MGE-derived interneurons (L) Sholl analysis on DIV6 Rac1/3wt and Rac1/3dmut interneurons (n=34-37cells per condition). (M) Graph indicating neurites length of the interneurons. Student’s t test,**P<0.01, ***P < 0.001. Error bars indicate SEM. Scale bars represent 4,2,10 μm.

### Axonal transport and lysosomal positioning during migration of interneurons are affected upon ablation of Rac1/Rac3

Axon transport is critical for the axon outgrowth and homeostasis and a good indicator of proper transport is the motility of lysosomes along the axons (Maday et al., 2014). Lysosomes consist of two distinct pools, a highly motile subset moving bi-directionally and a rather immobile one, located in the perinuclear region around the microtubule-organizing center (MTOC) (Cabukusta and Neefjes, 2018).To investigate whether the absence of Rac1/Rac3 in CINs affect lysosome transportation and motility, we isolated MGE-derived interneurons from Rac1/Rac3wt and Rac1/Rac3dmut embryos and cultured them for 3DIV. We visualized lysosomes and microtubules and recorded the axonal transport of lysosomes in growing axons by time lapse confocal microscopy (Fig. 4A, B). For each axon, we generated kymographs to analyze lysosomal locomotion (Fig 4C). First, we quantified the lysosomes detected per axon in both genotypes and categorized them according to the direction of movement in four groups: moving anterogradely, moving retrogradely, moving in alternating directions (reversal) and immobile/stationary lysosomes (Fig. 4D). Regarding the number of lysosomes per axon, there was no difference between genotypes. Interestingly, although there was no significant difference in the number of stationary, or anterogradely-moving lysosomes per interneuron axon, we observed a significant reduction in the number of retrogradely moving lysosomes and an increase in the number of lysosomes moving in alternating directions (reversal) in double mutant compare to control CINs (Fig. 4D).

**Fig. 4.**
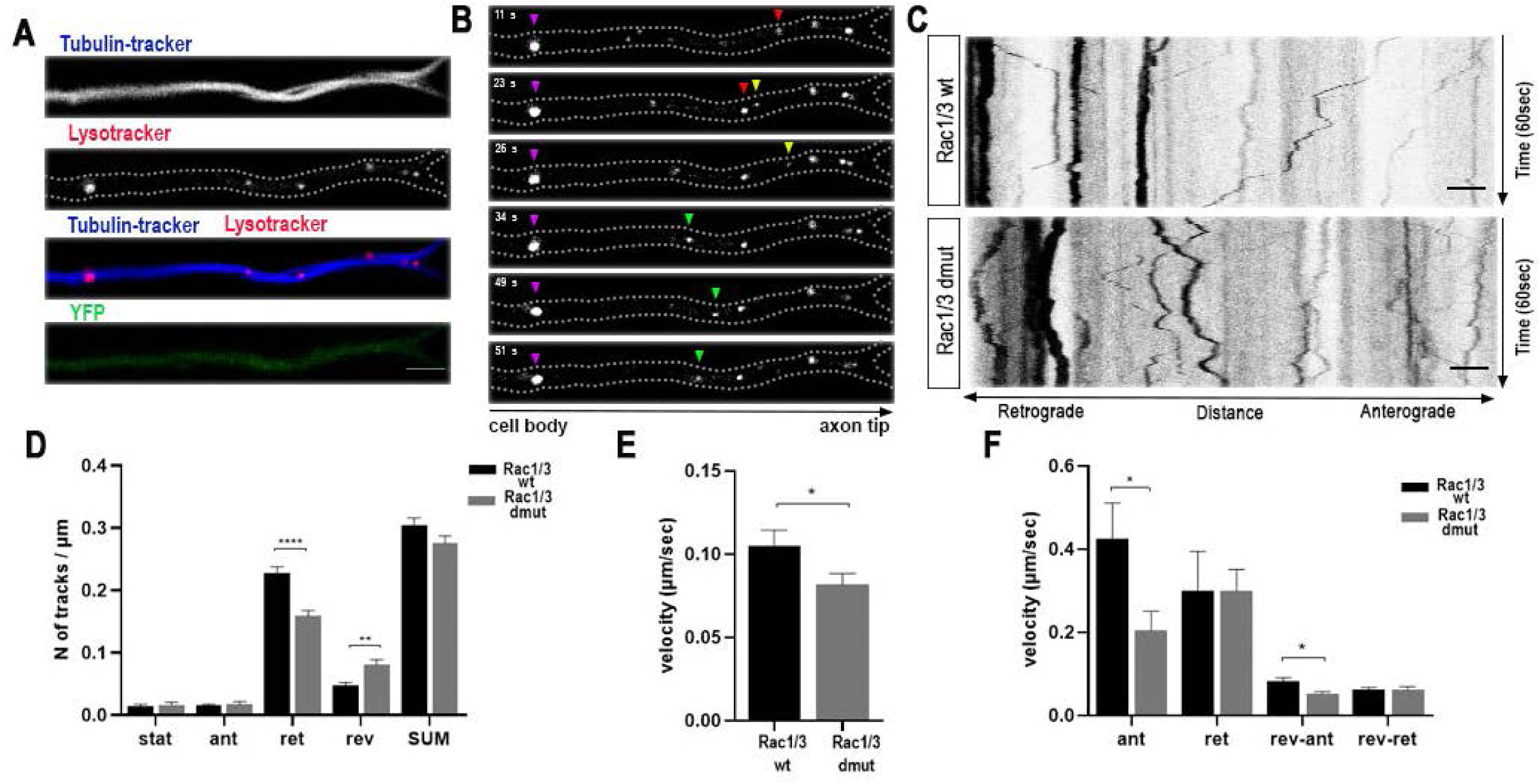
Lysosomal motility is impaired upon ablation of Rac1 and Rac3 in cortical interneurons. (A) Representative images of the axon of a YFP-positive (green) MGE-derived interneuron from E14.5 Rac1/3wt embryos, cultured 3DIV. Microtubules are stained with tubulin tracker (blue), lysosomes with LysoTracker (red). (B) Time-lapse series of the principal neurite demonstrating lysosomes moving in different directions. Purple arrowhead indicates a stationary lysosome, yellow arrowhead an anterogradely moving lysosome, red arrowhead a retrogradely and green arrowhead a lysosome moving in alternating directions. (C) Representative kymographs of lysosomes in the principal neurite segment of a Rac1/3wt and a Rac1/3dmut interneuron at 3DIV. (D) Histogram showing the densities of lysosomes along the principal neurite of Rac1/Rac3wt and Rac1/Rac3dmut interneurons per category (stationary, anterograde, retrograde, reversal). Lysosomal tracks are calculated as the number of tracks per axon length. (E) Histogram of the lysosomal velocities in μm/sec of Rac1/3wt and Rac1/3dmut interneurons. (F) Histogram of the lysosomal velocities in μm/sec of Rac1/3wt and Rac1/3dmut interneurons per category according to their direction of movement. (n=75-79 cells per genotype, from 4 independent experiments). Multiple t test, *P < 0.05, **P<0.01, ****P < 0.0001. Error bars indicate SEM. Scale bars represent 5 μm.

Regarding the velocity of mobile lysosomes, it was significantly reduced in Rac1/Rac3dmut compare to wt interneurons (Fig 4E). We further quantified the velocity of lysosomes according to their directionality. Except for the anterogradely and the retrogradely moving lysosomes, we divided also the group of the lysosomes with alternating directions into lysosomes with resultant anterograde or retrograde direction. There was a significant reduction in the lysosomes moving anterogradely and in the lysosomes with resultant anterograde direction of movement, while there was no significant alteration in the velocity of the retrogradely moving lysosomes (Fig 4F). Thus, by examining lysosomal locomotion, we concluded that axon transport was affected in Rac1/Rac3dmut interneurons, which could further affect their axonal outgrowth. Immunolabelling for LAMP-1, a marker of late endosomes/lysosomes revealed that in the migrating interneurons lysosomes appeared in the perinuclear region, at the axon initial segment as well as in the swelling formation (Fig S3). Therefore, to monitor the lysosomal positioning during a migratory cycle, we performed live imaging in interneurons migrating from MGE explants by using fluorescent probes to visualize the nucleus and lysosomes (sup video). Interestingly, while in Rac1/Rac3wt interneurons the lysosomal pool located in the perinuclear region moved forward, into the swelling, prior to the nuclear translocation, in the Rac1/Rac3dmut interneurons lysosomes were moving anterogradely simultaneously with the nucleus. This observation indicated that upon ablation of Rac1 and Rac3, the resulted defects in cytoskeletal organization affect also the proper positioning of lysosomes during the migratory cycle.

### Altered expression of genes in the medial ganglionic eminence upon ablation of Rac1/Rac3

In order to better understand the migratory defects we observe in CINs in the absence of Rac1/Rac3, we wanted to identify changes in gene expression levels, using RNA-sequencing of MGEs from both wt and dmut embryos. Setting a minimum average 2fold difference of expression as a criterion, we generated a list of genes (N=364) with altered expression; 224 of these genes exhibited decreased expression vs 140 genes with increased levels (fig. 5A-C). Upon Gene Set Enrichment Analysis, we focused on genes with products that are implicated in neuronal differentiation, cell cycle progression in neuronal progenitors or other cell types, genes involved in tumorigenesis and neurodevelopmental disorders, or genes associated with Rho signaling and related to cytoskeletal organization (table 1). To validate the expression effects of Rac1/Rac3 ablation for the majority of those genes, we employed quantitative RT-PCR (qPCR) with mRNA extracted from both, Rac1/Rac3wt and Rac1/Rac3dmut MGEs (fig. 5D).

**Fig. 5.**
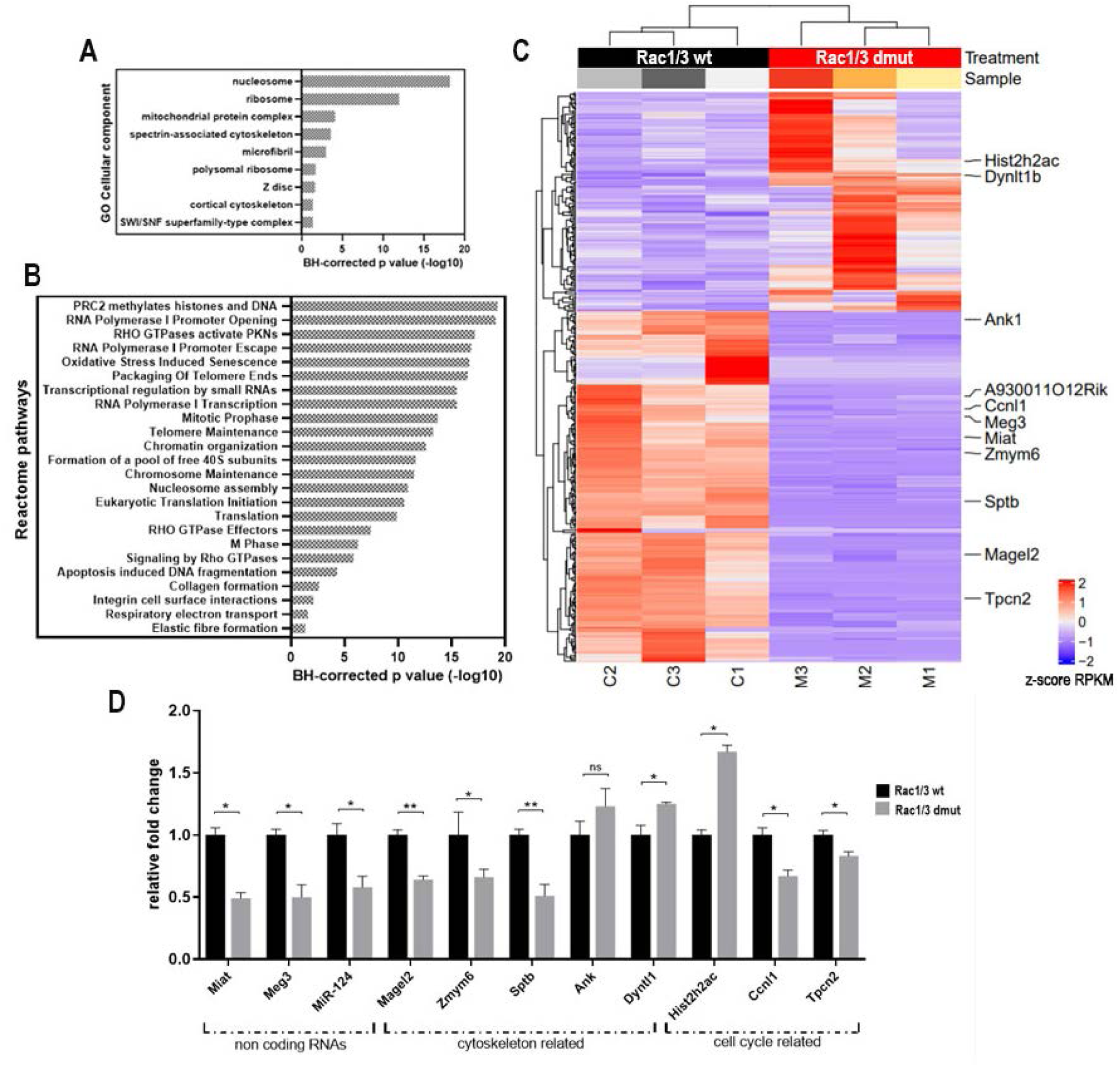
Altered gene expression in Rac1/3dmut MGEs. (A) Enriched GO terms for cellular components and (B) reactome pathways of genes with altered expression in Rac1/3dmut E13.5 MGEs acquired from RNAseq analysis. (C) Heat map of genes acquired from RNAseq analysis. (D) qRT-PCR of selected genes normalized to control (Rac1/3wt) and to Gapdh (n=3 samples per genotype). Student’s t test, *P<0.05, **P<0.01. Error bars indicate SEM.

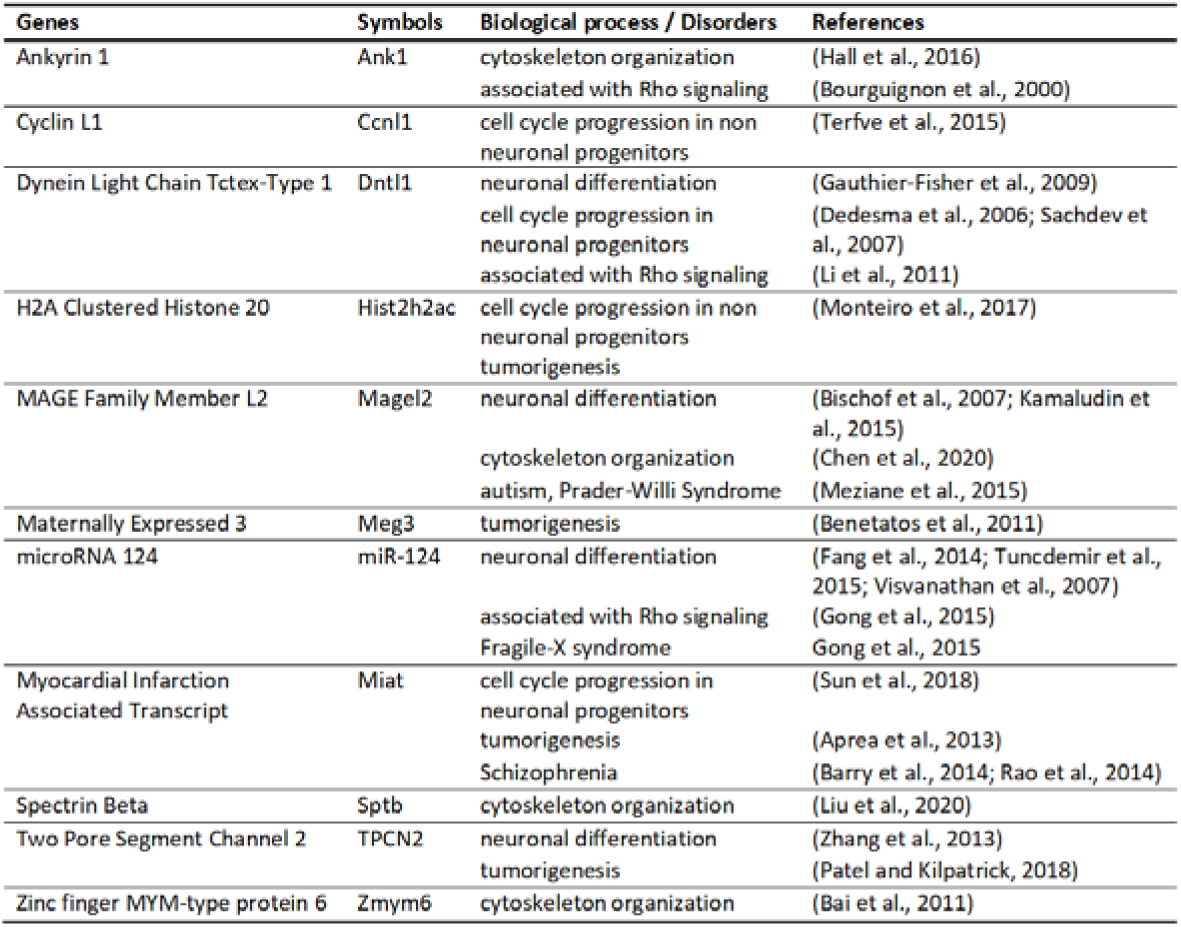

### Expression and distribution of TPC2 in interneurons is affected upon ablation of Rac1/Rac3 GTPases

Among the list of differentially expressed genes we identified (see table), we choose to further investigate the voltage-gated ion channel Two-Pore Channel 2 (TPCN2 gene encoding TPC2). TPC2 is mainly located in late endosomes/lysosomes and it mediates Ca^+2^ release upon activation by the second messenger nicotinic acid adenine dinucleotide phosphate (NAADP) and other molecules (Calcraft et al., 2009; Jha et al., 2014; Lee et al., 2016; Ogunbayo et al., 2018; Ruas et al., 2010; Zong et al., 2009). Interestingly, TPC2 is implicated in in cancer metastasis (Nguyen et al., 2017). Our RNAseq data revealed that the mRNA levels of the TPC2 gene were decreased in Rac1/Rac3dmut interneurons. To test the respective protein levels, we cultured for 3DIV MGE-derived interneurons from Rac1/Rac3dmut and control embryos, and then immunostained for YFP, TPC2 and LAMP-1, in order to mark late lysosomes where TPC2 is highly expressed (Fig. 6A-B’’). Measuring both fluorescence area (Fig. 6C) and intensity (Fig. 6D), we concluded that the levels of the TPC2 protein were significantly decreased in Rac1/Rac3d mut cells compare to control ones. To assess the TPC2 subcellular localization, we quantified its mean intensity across the axon, in standard distances (Fig. 6E). We noticed that in the first part of the axon, approximately in the first 7μm, there was no significant difference in expression levels, while there was a significant reduction in the intensity of the TPC2 in the rest of the neurite until the neurite tip in Rac1/Rac3dmut interneurons. We also noticed a significant increase in the perinuclear region of the TPC2 in Rac1/Rac3dmut compared to wt cells. Together these data demonstrate that the distribution of the TPC2 in the Rac1/Rac3dmut interneurons was altered. LAMP-1 distribution was in accordance with the pattern described for TPC2 (Fig. 6F). In summary, upon ablation of Rac1 and Rac3, the TPC2 levels were decreased and its subcellular localization was affected.

**Fig. 6.**
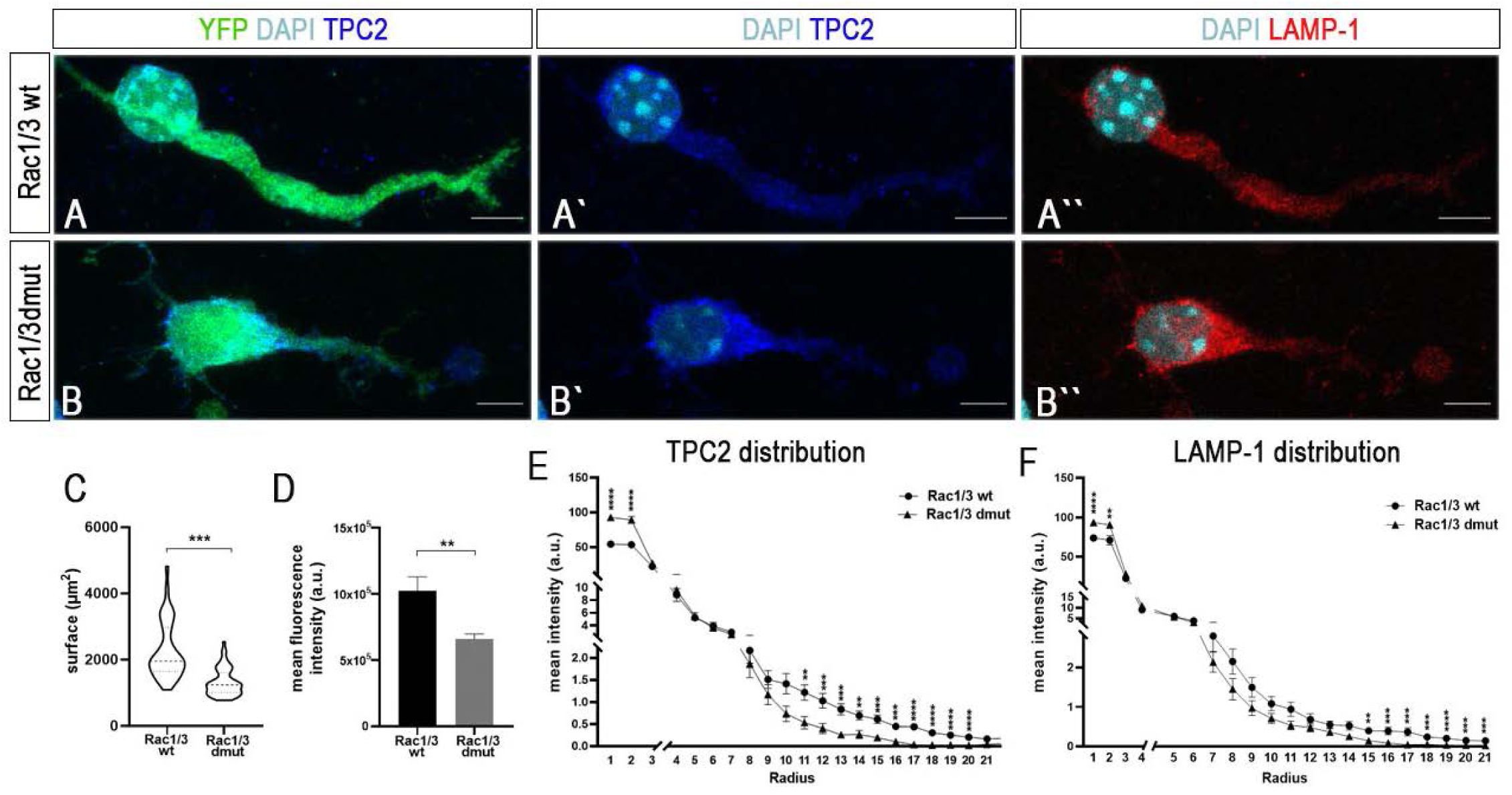
Depletion of Rac1 and Rac3 impairs the intensity and the subcellular distribution of TPC2 in MGE-derived interneurons. (A-B’’) Immunostaining of ΤPC2 (blue), YFP (green), late lysosomes (LAMP-1 in red) in cultured MGE-derived interneurons from E14.5 Rac1/3wt (A,À’) and Rac1/3dmut (B,B’’) embryos, 3DIV. The nucleus is labeled with DAPI (cyan). (C) Graph showing the surface, in μm^2^, occupied by TPC2. (D) Histogram indicating the intensity of TPC2 in Rac1/3wt and Rac1/3dmut interneurons. Graph showing the distribution of TPC2 (E) and of LAMP-1 (F) in Rac1/3wt and Rac1/3dmut interneurons measured in concentric circles with standard radii starting from the center of the nucleus. Student’s t test (C,D), multiple t test (E,F), *P<0.05, **P<0.01 ***P<0.001, ****P < 0.0001,. Error bars indicate SEM. Scale bars represent 5 μm.

### Pharmacological inhibition of TPC2 negatively affects axon outgrowth and migration of MGE-derived interneurons

We demonstrated that axon outgrowth is severely disturbed in Rac1/Rac3dmut interneurons (fig 3). Interestingly, a role of TPC2 in regulating caudal primary motor neuron axon extension has been recently uncovered in zebrafish (Guo et al., 2020). Moreover, recent studies point to role for TPC2 in calcineurin activation, which in turn activates Rac GTPases (Davis et al., 2020; Pandey et al., 2009). Therefore, we wanted to test whether TPC2 had a role in axon outgrowth of interneurons, similar to that of Rac1/Rac3, by examining the impact of its loss of function. TPC2-mediated Ca^+2^ release was inhibited pharmacologically by transned-19, a known NAADP antagonist (Guo et al., 2020; Kelu et al., 2015; Nguyen et al., 2017). We cultured MGE-derived interneurons from E14.5 Rac1/Rac3wt embryos and monitored the level of the axon extension upon treatment with trans-Ned 19. At DIV1 after plating, we treated the cells with trans-Ned 19 in two concentrations (20μΜ and 100μΜ) for 24hrs (Foster et al., 2019). We used as control the untreated cells and also interneurons treated with DMSO, the diluting reagent for trans-Ned 19. We performed immunostaining for YFP, F-actin to visualize the growth cones, LAMP-1 for the late lysosomes and DAPI to stain the nuclei (Fig. 7A-E’). We measured the length of the principal neurite, which will become the axon, in each condition (Fig. 7F). In the lower concentration of the inhibitor, we observed a significant reduction in axon length compared to the untreated and the DMSO treated cells (27%). In interneurons treated with the higher dose of the trans-Ned 19, the reduction in the axon length was even higher (46%). We also noticed an alteration in the width of the axon (data not shown) especially in interneurons treated with the high dose. Thus, we quantified also the surface of the axon, in order to explore the possibility that the gain in the width might compensate for the loss in the length (Fig. 7G). We detected a tendency towards reduction in the axon surface of the treated cells in the low concentration and a significant reduction in the axon surface at the higher dose of the trans-Ned 19, reaching about 45%. The decrease in the length and the surface of the axon was so pronounced that it had a negative impact in the total cell surface of the interneurons when the cells were treated with the high concentration of trans-Ned 19 (Fig. 7H). In this condition the total cell surface was found decreased about 30%. These data indicate that the pharmacological inhibition of the TPC2 negatively affects axon extension in the wild type MGE-derived interneurons.

**Fig. 7.**
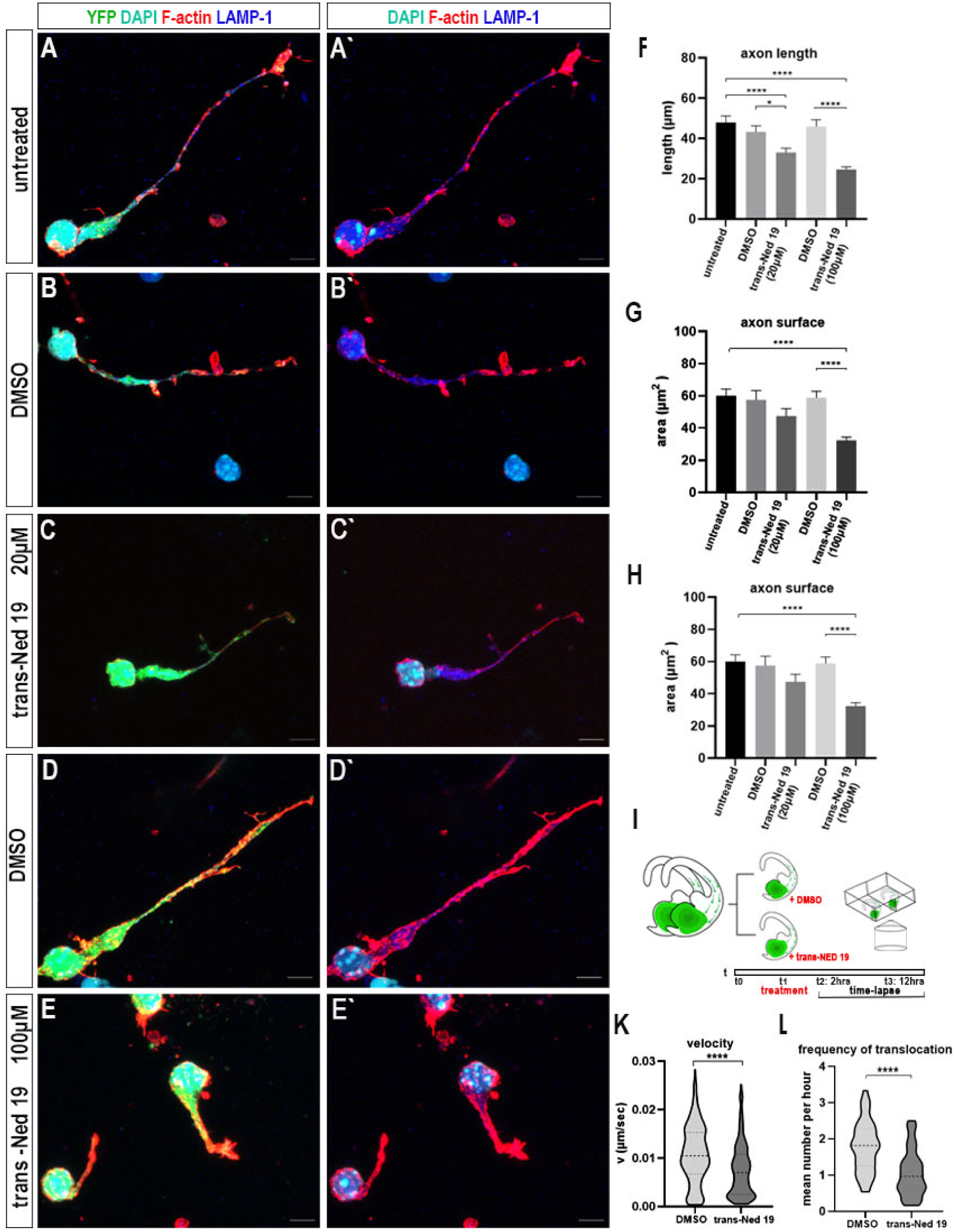
Pharmacological inhibition of TPC2 negatively affects axon outgrowth and migration of wild type interneurons. (A-E’) Immunostaining for F-actin (phalloidin, in red), YFP (green), late lysosomes (LAMP-1 in blue), in cultured MGE-derived interneurons from E14.5 Rac1/3wt embryos, 2 DIV. The nucleus is labeled with DAPI (cyan). Interneurons were untreated (A-A’) or were treated with DMSO (B-B’, D-D’) or trans-Ned 19 (C-C’, E-E’) at 20μΜ and 100μM concentrations respectively. Quantification of the axon length (F), axon surface (G) and the total cell surface (H) (n=42-50 cells per condition, from 3 independent experiments). One-way ANOVA Tukey’s multiple comparisons test, *P<0.05, ***P<0.001, ****P < 0.0001. (I) Schematic representation of the treatment overview. E14.5 brain slices treated with 200μΜ trans-Ned 19 and DMSO were recorded simultaneously for ∼8hrs. (time-lapse images acquired every 6min) Quantification (K) of the velocity of the migrating interneurons in presence of DMSO or trans-Ned 19 (number of embryos: 4, n=52-60 cells per condition) (mean±SEM 0.0108±0.00038 Rac1/3wt, 0.0075±0.00035 Rac1/3dmut), (L) the frequency of nuclear translocation of migrating interneurons (number of embryos: 4, n=45-49 cells per condition), (mean±SEM: 1.787±0.1038 Rac1/3wt, 1.062±0.0916 Rac1/3dmut), Student’s t test, ****P < 0.0001. Error bars indicate SEM. Scale bars represent 10 μm.

Upon ablation of Rac1 and Rac3, the rate of axon outgrowth was significantly altered particularly in the period of 2-4 hours (Fig. 3). Therefore, we asked whether the inhibition of TPC2 might have a similar impact. We performed a protocol where Rac1/Rac3wt MGE-derived interneurons were treated with the high dose of trans-Ned 19 one hour after plating. We stopped the treatment at different time points (2hrs, 4hrs and 12hrs later, Fig. S4A). We stained the interneurons with phalloidin, YFP and LAMP-1 and we measured the axon length and the axon surface in each time point as above (Fig. S4B-J’). After 2hrs of treatment, there was a tendency towards reduction in the axon length and the axon surface at the trans-Ned 19-treated interneurons compared to the untreated and the DMSO-treated ones (Fig. S4K). After 4hrs of treatment, there was a significant reduction in both axon length and surface. The length in the trans-Ned 19 treated cells was decreased about 45% compared to the DMSO-treated interneurons and the axon surface about 50% (Fig. S4L). In the next time point, at 12hrs, this significant reduction in the length and the axon surface was maintained. The axon surface was reduced about 35% at this time point in the treated interneurons indicating that the most pronounced difference in the axon outgrowth was in the period from 2 to 4 hours after treatment, corresponding to 3-5 hrs after plating. Together, these data indicate that the pharmacological inhibition of the TPC2, negatively affects the initiation of the axon outgrowth, as well as its maintenance.

Lastly, we aimed to investigate whether the process of migration might be impaired upon inhibition of TPC2. In order to answer this question we prepared E14.5 organotypic brain slices from Rac1/Rac3wt embryos. We treated them with trans-Ned 19 following a long-term treatment protocol (250μΜ of the inhibitor, for 12hrs). We treated the slices for 4hrs and then performed live imaging in the treatment solution for 8 hrs, simultaneously for the DMSO and the trans-Ned 19-treated brain slices (Fig. 7I). We recorded the interneurons migrating from the subpallium to the cortex and quantified their mean velocity (Fig. 7K). We found that there was 30% reduction and a significant decrease in the frequency of translocation (Fig. 7L), suggesting that the pharmacological inhibition of the TPC2 negatively affected the migration of interneurons.

Collectively, our data support the idea that reduction of TPC2 upon ablation of Rac1 and Rac3 could contribute to the phenotype observed regarding the initiation of axon outgrowth, axon extension and interneuron migration.

## Discussion

In this manuscript, we present evidence that certain cellular parameters of CIN migratory behavior, such as: centrosome, Golgi and lysosome positioning, are perturbed upon Rac1/Rac3 ablation. In addition, we show that axon growth and in particular, axon initiation, are severely affected. We also provide evidence that the expression and localization of the two-pore channel 2 (TPC2), a voltage-gated ion channel mediating Ca^2+^ release, are altered in Rac1/Rac3 mutant CINs. Interestingly, pharmacological inhibition of TPC2 negatively affects axonal growth and migration of interneurons *in vitro*. Taken together, our data suggest that TPC2 contributes to the severe phenotypes, concerning axon growth initiation, extension and migration that are observed in CINs, which lack Rac1 and Rac3 expression.

### Ablation of Rac1/Rac3 GTPases results in impairments in the migration of cortical interneurons

Here, we describe how the loss of Rac1/Rac3 affects the three-step “movement cycle” of migrating MGE-derived interneurons. The velocity of the migration of Rac1/Rac3dmut interneurons is significantly decreased. The average duration of the cycle is increased, while the frequency and the amplitude of nuclear translocation are reduced.

In migrating neurons, actomyosin contractility plays a fundamental role in driving nuclear translocation (Baudoin et al., 2007; Bellion, 2005; Martini and Valdeolmillos, 2010; Schaar and McConnell, 2005b). Phosphorylated myosin II, via RhoA and Rho-kinase signaling, that accumulates at the rear of MGE-derived interneurons, promotes the forward movement of the nucleus (Bellion, 2005; Laguesse et al., 2012; Peyre et al., 2015). In CINs upon ablation of Rac1/Rac3, the levels of RhoA and phosphorylated-myosin II are reduced, indicating an impairment in the actomyosin contractility of the cell, likely resulting in abnormal nucleokinesis and decreased velocity in migrating neurons.

In our mutant CINs the characteristic swelling formation that initiates the migratory cycle is formed closer to the nucleus, and might contribute to the abnormal localization of the centrosome in short distance from the nucleus, as well as, the slower nuclear translocation. The above defects in CIN nucleokinesis observed in our Rac1/Rac3dmut neurons could be further explained by alterations in microtubule dynamics in the perinuclear region (Fig. S2). We have previously observed a reduction in ac-tubulin in Rac1/Rac3dmut interneurons (Tivodar et al., 2015). These observations are in agreement with other reports showing that instability of the microtubules (as well as excessive stability) affects the nuclear translocation and the distance of the centrosome from the nucleus (Garcez et al., 2015; Tanaka et al., 2004; Umeshima et al., 2007; Valiente et al., 2010). MGE-derived interneurons treated with nocodazole, which destabilizes microtubules, exhibit shortened leading processes and decreased migration speed (Baudoin et al., 2007). This phenotype resembles that of Rac1/Rac3dmut interneurons we describe here. In addition, deletion of doublecortin, which stabilizes microtubules, results in interneurons with branching phenotypes and abnormalities in nuclear translocation and swelling (Kappeler et al., 2006). Accordingly, doublecortin-deficient neurons migrating in the rostral migratory stream in the adult brain, also exhibit impairments in nuclear translocation, however, unlike Rac1/Rac3dmut interneurons, no defect in centrosome positioning is observed (Koizumi et al., 2006). Finally, in glial-guided migration of cerebellar granule neurons (CGNs), myosin II and actin cytoskeleton play a role in the harmonized translocation of the centrosome and soma (Solecki et al., 2009).

Our analysis also showed that the formation of the swelling persists for a longer period, suggesting that RhoGTPase activity might have a role in its modulation. In agreement with this, in excitatory cortical neurons, it has been shown that the Rac1 interacting protein POSH is required to properly localize activated Rac1 to the cytoplasmic dilation, a cellular feature similar to the swelling formation observed in migrating CINs (Yang et al., 2012).

Finally, the Golgi Complex also moves to the swelling prior to nuclear translocation in CINs (Baudoin et al., 2007; Yanagida et al., 2012). Upon ablation of Rac1/Rac3, MGE-derived interneurons display abnormal Golgi positioning, which is now localized closer to the nucleus. As discussed for the centrosome above, it has been shown that in migrating CGNs, F-actin and Myosin II are necessary for the Golgi translocation (Trivedi et al., 2014). In addition, in hippocampal neurons, dendritic Golgi positioning is influenced by microtubule dynamics (Meseke et al., 2013a). Our data are also in agreement with the described phenotype in hippocampal neurons, where dominant negative Rac1 and Cdc42 alters dendritic Golgi positioning (Meseke et al., 2013b). Additional organelles, such as mitochondria are also motile during CIN migration (Lin-Hendel et al., 2016). Here, we present new evidence that lysosome positioning changes during the three-step cyclic migration of MGE-derived interneurons and that in the absence of Rac1/Rac3 possibly due to impairments in the cytoskeleton organization this positioning is altered.

### Defects in cytoskeleton organization affect the axon growth in Rac1/Rac3 deficient interneurons

Rho GTPases, including Rac1 and its effectors STEF, Tiam, FIR, DOCK7 are known to have profound effects in neurite initiation and axon outgrowth (Gonzalez-Billault et al., 2012; Govek et al., 2005; Kunda et al., 2001; Matsuo et al., 2002; Schelski and Bradke, 2017a; Tanaka et al., 2004; Watabe-uchida et al., 2006). Conditional ablation of Rac1 from different neurons revealed a requirement of this GTPase in axon growth and guidance (Hua et al., 2015). Moreover, in CGNs that do not express other Rac isoforms, ablation of Rac1 resulted in impairments in neuronal migration and axon formation (Tahirovic et al., 2010).

Here, we provide evidence that the ablation of Rac1/Rac3 from CINs affects axon initiation, as well as, the rate of axonogenesis. We reveal that in wt CINs the rate of axon outgrowth is increased within the first 5 hrs after plating. In contrast, Rac1/Rac3dmut interneurons show a significant decrease in the rate of neurite outgrowth during the same time frame, indicating a defect in axon initiation. Axon elongation was also impaired in Rac1/Rac3dmut interneurons, as they have shorter axons compare to control ones, after 6DIV. Additionally, Rac1/Rac3dmut interneurons exhibit an over-branched phenotype possibly as a result of the reduced levels of ac-tubulin, an observation that is in agreement with previous work in hippocampal neurons (Wei et al., 2018).

Actin-mediated protrusions and microtubule-mediated forces present in the growth cone, have a fundamental role in axonal extension (Coles and Bradke, 2015). Rac1 is a known regulator of actin dynamics via the WAVE complex, thus by increasing the actin filaments providing the forces needed for leading edge extension (Schelski and Bradke, 2017b; Tahirovic et al., 2010). In addition, Rac1 facilitates microtubule polymerization by inhibiting Stathmin (Watabe-Uchida et al., 2006; Wittmann et al., 2004). Moreover, activation of Rac1 is coupled to microtubules by Rac1 GEF triple functional domain (Trio) and MAP1B interactions, promoting axon growth (Montenegro-Venegas et al., 2010; Schelski and Bradke, 2017b; Van Haren et al., 2014). Ablation of Rac1/Rac3 from CINs affected severely the morphology of the growth cone, which could result in certain alterations of the described forces, leading in the defective axonogenesis displayed in our model.

A balanced regulation of Rac1 also promotes neurite extension by stabilizing the paxillin-containing contact points (Woo and Gomez, 2006). Its loss in our model could be an additional cause resulting in defective axonal outgrowth. A p21-activated kinase 1-interacting exchange factor (PIX)/ G-protein-coupled receptor (GPCR) kinase interacting protein1 (GIT) network has been identified to play important roles in the neuritogenesis and maturation of MGE-derived interneurons (Franchi et al., 2016). Interestingly, in a neuronal cell line Rac1 and Rac3 were found to modulate differentially GIT1-paxillin signaling, resulting in opposing effects on cell morphology and neuronal differentiation (Hajdo-Milasinovic et al., 2009). Future work examining whether this signaling pathway might affect axonal outgrowth initiation in interneurons and whether distinct roles of Rac1/Rac3 could be identified, may provide new insights in the role of GTPases in neuronal development.

### Defects in axonal transport in Rac1/Rac3 deficient interneurons

Axonal transport is a fundamental process underlying axon growth (Guillaud et al., 2020; Martenson et al., 1993). Lysosomes transport materials in axons over long distances, a process necessary for the growth of the developing axon (De Pace et al., 2020). Here we show that upon ablation of Rac1/Rac3, lysosomal motility is impaired in elongating axons of interneurons. Lysosomes move bidirectionally in a “stop and go” motion, with frequent pauses, coupled to kinesin and dynein motors (Farías et al., 2017; Guedes-Dias and Holzbaur, 2019; Hendricks et al., 2012; Pu et al., 2016). Several kinesins mediate anterograde lysosome translocation (Kanai et al., 2000; Nakata et al., 2011; Xia et al., 1998). We show that in Rac1/Rac3dmut interneurons the number of the retrogradely moving lysosomes per axon is decreased. However, this lower number is in part compensated by more lysosomes moving in alternating directions that finally have a resultant retrograde flow (fig 4D). The velocity of the moving lysosomes is reduced mainly in the ones translocated anterogradely (fig. 4F). Post translational modifications of tubulin are known to affect the kinesin motility (Guedes-Dias and Holzbaur, 2019; Schelski and Bradke, 2017a). Kinesin-1 moves preferentially along stable acetylated or detyrosinated microtubule tracks (Cai et al., 2009). KIF5C, a protein of kinesin-1 motor, localizes in stable detyrosinated mictotubules, while kinesin transport is promoted by acetylation of tubulin (Dunn et al., 2008; Reed et al., 2006). Alterations in acetylation levels lead to a reduction in velocity in organelles and vesicles (Dompierre et al., 2007; Godena et al., 2014; Morelli et al., 2018). The reduction of ac-tubulin that the Rac1/Rac3dmut interneurons exhibit, might be the reason for the slower anterograde lysosomal transport. It has been demonstrated that kinesin-1 moves faster on acetylated or GTP-bound stable microtubules (Pu et al., 2016). Additionally, motility of intracellular organelles including lysosomes, is regulated by Ca^+2^ (Li et al., 2016). Thus, alteration in Ca^+2^ transients might also result in abnormal lysosomal motility. Future studies may address the question of whether the defective lysosomal motility has also consequences on the cargoes involved in this process.

### Two pore channel expression in interneurons is altered upon ablation of Rac1/Rac3 GTPases

The ablation of Rac1/Rac3 from MGEs at E13.5 results in alterations in gene expression (Fig 5). Genes associated in previous studies with the phenotype observed in Rac1/Rac3 deficient mice were selected according to the criteria described in table 1 and the expression levels were validated. Among them we noticed the Two-pore channel 2, a voltage-gated ion channel of late endosomes and lysosomes (Calcraft et al., 2009; Ruas et al., 2010). Out of the three existing channels, only TPCN1 and TPC2 are expressed in humans and mice (Brailoiu et al., 2010; Calcraft et al., 2009). TPC2 mediates intracellular Ca^+2^-release from lysosomes after activation with low nanomolar concentrations of NAADP and other molecules such as phosphatidylinositol 3,5-bisphosphate (PI(3,5)P2), Mg^+2^ and the mitogen activated protein kinases (MAPKs) c-jun N-terminal kinase (JNK), P38 and the mechanistic target of rapamycin complex 1 (mTORC1). The Ca^+2^-release is lysosome-specific (Calcraft et al., 2009; Jha et al., 2014; Ogunbayo et al., 2018; Zong et al., 2009). More importantly, NAADP Ca^+2^ release trigger further Ca^+2^ signals via the endoplasmic reticulum (Calcraft et al., 2009; Galione, 2015; Ruas et al., 2010). In a proteomic characterization of the human TPC interactome Rac1 has been identified as a protein fulfilling the criteria of a TPC2 interactor (Lin-Moshier et al., 2014). This interaction was found with mass spectrometry and was nοt validated with co-immunoprecipitation as currently there was no *in vivo* evidence of localization of Rac1, among other proteins, with acidic stores. Our data provide an additional indication of an existing association of Rac1 and/or Rac3 with TPC2.

A role of Ca^+2^ signaling, NAADP and TPC2 in neuronal differentiation have been described. Spontaneous Ca^+2^ signaling was identified as an intrinsic characteristic of differentiating neurosphere precursors derived from embryonic striatal cells (Ciccolini et al., 2003). Later on, liposomal delivery of NAADP into PC12 cells was shown to mediate Ca^+2^ release from acidic stores and this release was sufficient to drive differentiation of the cells to a neuronal-like phenotype (Brailoiu et al., 2006). TPC2 knockdown in ΕS cells trap them in the neural progenitor stage by facilitating their differentiation into neural progenitors which fail to progress into the final neuronal stage (Zhang et al., 2013).

Upon ablation of Rac1 and both Rac1/Rac3 we have demonstrated a defect in the cell cycle of the MGE-derived progenitors. Rac1 is necessary for their transition from G1 to S phase and the deficient progenitors exhibit a prolonged G1 Phase (Tivodar et al., 2015; Vidaki et al., 2012). A working hypothesis might be that the reduced levels of TPC2 observed in the absence of Rac1/Rac3 reflect the different stage of the progenitors in the VZ and the SVZ of the MGE of the dmut mice.

### Two-pore channel 2 affects axon growth and migration in CINs

Ca^+2^ signaling regulates several cellular processes in neurons such as axon guidance, growth cone motility, neurite extension and branching, as well as synaptogenesis (Gomez and Zheng, 2006; Zheng and Poo, 2007). In Xenopus spinal neurons and also in cortical neurons, the rate of axon growth was inversely correlated with the frequencies of spontaneous Ca^+2^ transients (Gomez and Spitzer, 1999; Tang et al., 2003). Plasma membrane influx along with intracellular stores mobilization result in Ca^+2^ concentration alterations. NAADP-induced Ca^+2^ release from intracellular stores were found to potentiate neurite outgrowth in rat cortical neurons (Brailoiu et al., 2005). More recently, TPC2-mediated Ca^+2^ signaling has been implicated in axon extension of caudal primary motor neuron in zebrafish, through the use of the NAADP receptor antagonist trans-ned-19, as well as genetic silencing strategies (Guo et al., 2020). In agreement with these results, we show here that *in vitro* pharmacological inhibition of TPC2 function with trans-Ned 19 disturbs axon outgrowth initiation and axon extension in CINs. Thus, the reduced levels of TPC2 observed in Rac1/Rac3dmut CINs might partially contribute to the described decreased axon outgrowth rate.

Finally, we demonstrate a requirement of TPC2 in interneuron migration. *Ex vivo*, pharmacological inhibition of TPC2 function with trans-Ned 19 severely impaired the migration of interneurons from the ventral telencephalon towards the developing cortex. Interestingly, TPC2 was also implicated in metastatic invasion which also requires active migration. Pharmacological inhibition or silencing of TPC2 *in vitro* reduced adhesion, leading edge formation and migration of cancer cells, while metastasis *in vivo* was also reduced (Nguyen et al., 2017). The inhibition of TPC2 function was shown to disturb trafficking of β1-integrin, a protein that is involved in tumor migration (Nguyen et al., 2017). Disturbances in Ca^+2^ homeostasis are also probable (Alharbi and Parrington, 2019). We propose that the defective migration described in Rac1/Rac3dmut CINs could be in part due to the reduction of TPC2.

Ca^+2^ release from endoplasmic reticulum mediated by the ryanodine-and IP3-sensitive channels regulates motility and directionality of migrating neurons (Guan et al., 2007; Horigane et al., 2019; Hutchins et al., 2011; Liu et al., 2008). Thus, a working hypothesis could be that altered Ca^+2^ released from intracellular stores upon inhibition of TPC2 might have a negative impact in the migration of interneurons.

A crosstalk between Ca^+2^, small Rho GTPases, their effectors and cytoskeleton remodeling exists, affecting migration and axon growth described in different neuronal types such as CGN and cortical neurons (Gomez and Zheng, 2006; Kholmanskikh et al., 2006; Nicole et al., 2018). In Rac1/Rac3dmut interneurons the cytoskeleton remodeling primarily results from the ablation of the two Racs; however, this existing crosstalk might support the hypothesis that altered Ca^+2^ transients upon reduction of TPC2 might contribute to the defective phenotype.

Changes in intracellular Ca^+2^ in growth cones arise in response to Ca^+2^ influx either through plasma membrane channels or release from intracellular stores (Gomez and Zheng, 2006). It has been shown that calcium release through IP3 receptors regulates migration in Gonadotropin-releasing hormone (GnRH)-expressing neurons, through a signaling pathway involving the calcium sensor, calcium/calmodulin protein kinase kinase, AMP-activated kinase, and RhoA/ROCK, stimulating actin flow at the leading process (Hutchins et al., 2013). TPC2 localization in the growth cones might contribute to the intracellular Ca^+2^ changes and the regulation of the growth cone. Moreover, actomyosin contraction at the cell rear providing forces for nucleokinesis of the tangentially migrating interneurons requires spontaneous fast intracellular calcium transients while myosin light chain kinase (MLCK) which promotes actomyosin contractility is a downstream molecule of calcium/calmodulin complex (Martini and Valdeolmillos, 2010). The localization of the rather immobile pool of lysosomes in the perinuclear region of the interneurons raise the possibility of local Ca^+2^transients required for the actomyosin contraction created by lysosomal Ca^+2^ release through TPC2. In accordance with this hypothesis the reduction of the TPC2 found in Rac1/Rac3dmut interneurons would have complementary effects, resulting in the observed abnormal nucleokinesis.

Taken together our data underscore the importance of Rac1/3 in the early phases of axon growth and migration of the cell body in multiple cellular processes impacting the developmental journey of interneurons to the cortex. Moreover, the two-pore channel 2, a lysosomal calcium channel implicated in the migration of cancer cells is also shown to be involved in neuronal development, mainly in migration and axonogenesis of interneurons.

## Material and methods

### Animals

All animals were kept at temperature-controlled conditions on a 12 h light/dark cycle, fed by standard chow diet and water ad libitum provided within the animal facility of the Institute of Molecular Biology and Biotechnology (IMBB)-Foundation for Research and Technology Hellas (FORTH) (license nos. EL91-BIObr-01 and ΕL91-BIOexp-02). All animal experiments complied with the ARRIVE and NC3Rs guidelines to improve laboratory animal welfare and conformed with all regulations and standards outlined in the Presidential Decree 56/ 30.04.2013 (Greek Law) in accordance with the EU directives and regulations (2010/63/EU and L 276/33/20.10.2010) and to the U.K. Animals (Scientific Procedures) Act, 1986 and associated guidelines, equivalent to NIH standards. Animals carrying a floxed allele of Rac1 (Rac1^fl/fl^;Nkx2.1^+/Cre^) were previously described (Vidaki et al., 2012) and will be referred as Rac1mut. Rac1 and Rac3 double-mutant animals were also previously described (Tivodar et al., 2015). Briefly, the Rac1 ^+/fl^;Nkx2.1^+/Cre^ line was crossed with the Rac3 ^KO^ line (Corbetta et al., 2005). In order to visualize Rac1/Rac3 mutant (and control) interneurons, the ROSA26^fl-STOP-fl-YFP^ allele was also inserted as an independent marker labeling the cells via the expression of yellow fluorescent protein (YFP) (Srinivas et al., 2001). Rac1^+/fl^;Rac3^+/+^;Nkx2.1^Tg(Cre);^R26RYFP^+/−^ and Rac1^fl/fl^;Rac3^−/−^; Nkx2.1^Tg(Cre);^ R26R-YFP^+/−^ animals will be referred as Rac1/Rac3 wt and Rac1/Rac3 dmut respectively, in Materials and Methods, text, figures, and legends.

### Culture of dissociated MGE-derived interneurons

MGEs were dissected from forebrains of E14.5 mouse embryos Rac1/Rac3 wt and Rac1/Rac3 dmut in ice cold HBSS/HEPES and treated with trypsin (0.05%, Gibco, Thermo Fisher Scientific, MA USA) for 10 min at 37 °C in 5% CO2. Subsequently, the cells were plated on coverslips coated with collagen (5 mg/mL) (Gibco) and cultured in Neurobasal/B27 medium, (2% B27 supplement, 0.5% glucose, 1X Glutamax, 1% Pen/Strep, in Neurobasal medium –phenol red, all Gibco) in a 5% CO2 humidified incubator at 37 °C as previously described (Vidaki et al., 2012). For long-term culture interneurons were plated on glass coverslips coated with matrigel matrix (Corning, Glendale, AZ, USA) (Franchi et al., 2018). For the recording of the initiation of axon outgrowth interneurons were embedded in Matrigel matrix on 35mm glass bottom culture dishes (ΜatTek Corporation, Ashland, MA) following the protocol previously described for MGE explants (Métin et al., 1997).

### MGE explants and organotypic embryonic brain slices cultures

Forebrains from E14.5 Rac1/Rac3 wt and Rac1/Rac3 dmut were dissected in ice-cold Leibovitz’s L15 medium (Gibco) and embedded in 4% low melting point agarose type VII (Sigma). Vibratome (Leica VT1000 S) 250μm thick coronal slices of mouse telencephalon were made and placed in 35 mm glass bottom culture dishes (ΜatTek Corporation) or (μ-slide, ibidi GmbH (FCA Gräfelfing, Germany) for live imaging experiments. Alternatively, MGEs were dissected out from the organotypic slices and were embedded in matrigel matrix on glass bottom culture dishes (Nunc). MGE explants from Rac1/Rac3 wt and Rac1/Rac3 dmut were cultured in Neurobasal/B27 medium for 1 DIV in a 5% CO2 humidified incubator at 37 °C and next prepared for live imaging acquisition. When telencephalic slices were prepared for focal electroporation the procedure was conducted in ice-cold Krebs solution (Stühmer et al., 2002).

### Focal electroporation on embryonic brain slices

Electroporation of MGEs on Rac1/Rac3 wt and Rac1/Rac3 dmut slices were performed as previously described (Stühmer et al., 2002). Plasmid DNA solution containing expression vector pCAGGs-IRES-RFP at a concentration of 1 µg/µL and fast green solution (stock 25 mg/mL) in a 1/10 dilution, were injected in the MGE of E14.5 brain slices placed on supporting membranes (Whatman nucleopore track-etched membranes, Sigma). Electroporation was conducted with platinum electrodes (ΒΤΧ Harvard Apparatus, Holliston, MA) P5, Sonidel) using ECM-830 BTX square wave electroporator (BTX Harvard Apparatus) (2 pulses of 125V of 5ms duration each at 500ms intervals). Electroporated slices were placed for 1 h in MEM medium (10% FBS, 0.5% glucose, 1% Pen/Strep, in MEM with glutamine, all Gibco) in a 5% CO2 humidified incubator at 37 °C and then incubated in Neurobasal/B27 medium for 18-24 h followed by the time lapse imaging (Denaxa et al., 2019).

### Pharmacological treatment

Trans ned-19 (Tocris Bioscience), the nicotinic acid adenine dinucleotide phosphate (NAADP) antagonist, was dissolved in DMSO and stored at-20°C. MGE-derived interneurons from E14.5 Rac1/Rac3 wt embryos were cultured in Neurobasal/B27 medium in a 5% CO2 humidified incubator at 37 °C and 24h after plating cells were treated with trans Ned-19 by exchanging the medium with one containing 25μΜ and 100μΜ trans ned-19 (Foster et al., 2019; Padamsey et al., 2017). Interneurons were treated for 24h. In a second set, cells were plated in Neurobasal/B27 medium and 1h later, they were treated, as above, with 100μΜ trans ned-19 for 24h. Control conditions included equal amounts of the DMSO solvent. Telencephalic brain slices from E14.5 Rac1/Rac3wt embryos, two per embryo were prepared, one treated with DMSO and one with trans-Ned 19, at 200μΜ concentration for 4h and then live imaging was performed in the treatment solution for additional 8h (Kelu et al., 2015; Nguyen et al., 2017). To manage simultaneous time lapse imaging of the trans Ned-19 treated brain slices and the DMSO treated slices, compartmentalized glass bottom slides were used (ibidi GmbH, Gräfelfing, Germany).

### Fluorescent dyes

MGE-derived interneurons from E14.5 Rac1/Rac3wt and Rac1/Rac3dmut embryos, at 4DIV with genotypes, were incubated for 1h in LysoTracker Red DND-99 reagent (Invitrogen) 75nM concentration and for 30 min in 1X tubulin tracker deep red reagent (Invitrogen) diluted in supplement-free medium according to the manufacturer’s instructions. MGE explants from E14.5 Rac1/Rac3wt and Rac1/Rac3dmut embryos, 24h after plating were incubated for 2h in LysoTracker Red DND-99 (75 nM) and for 45 min in Hoechst 33342 (Invitrogen) (0.25μg/ml).

### Immunostaining

For immunostaining in dissociated cells or MGE explants, interneurons grown in coated glass coverslips were fixed in 4% paraformaldehyde (PFA) or in 3.7% Formaldehyde, Methanol-free (for phalloidin staining) for 10 minutes and 15 minutes at room temperature (RT), respectively. Following fixation, cells were washed in PBS (3 washes, 10 min each for the cells and 20 min for the explants). Then the cells were permeabilized in PBS containing 0.1% Triton X-100 (PBST) for ∼15 min at RT and washed in PBS (3 washes, as described above). After permeabilization cells were incubated in 1% BSA in PBS for 1h at RT, while the MGE explants for blocked for 3h RT and subsequently incubated with primary antibodies, diluted in blocking solution, for 18h at 4°C. Next, cells were washed (3 times) and incubated with the secondary antibodies, diluted in blocking solution, for 2h at RT. At the end of this incubation step, the cells were rinsed with PBS and mounted, using MOWIOL (Calbiochem, Darmstadt, Germany). For immunostaining in embryonic brain slices, the slices were treated as the MGE explants described above with the difference that the blocking solution contained 3% FBS in PBST, and the antibodies were diluted in 1% FBS in RBST. The slices were incubated with primary antibodies for 18h at 4°C and for 18h at 4°C with secondary antibodies. Primary antibodies used: mouse monoclonal anti Acetylated-Tubulin (1:200; Sigma-Aldrich, Saint Louis, MI,USA), rabbit polyclonal anti-GFP (1:500; Minotech biotechnology, Heraklion, Greece), rat monoclonal anti-GFP (1:2000; Nacalai Tesque, Kyoto, Japan), mouse monoclonal anti-Golgin-97 (1:100; Thermo Fisher Scientific, MA USA), mouse monoclonal anti-Lamp-1 (1:50; Santa Cruz Biotechnology, California, USA), rabbit polyclonal anti-Pericentrin (1:500; Sigma-Aldrich), mouse monoclonal anti-Phospho-Myosin light chain 2 (Ser19) (1:200; Cell signaling, Beverly, MA, USA), anti-mouse Phospho RhoA, (NewEast Biosciences, PA), rabbit anti-RhoA (Cell Signalling Technology, Danvers, MA,USA), rabbit polyclonal anti TPCN2 (1:50; alomone labs, Jerusalem Biopark, Israel), mouse monoclonal anti-Tuj1 (1:1000; Covance, NJ, USA), rat anti-Tyrosinated Tubulin (1:100, Sigma-Aldrich). Fluorochome labeled secondary antibodies Alexa Fluor 488, 555 and 647 (1:800, Invitrogen, Carlsbad, CA, 1:800), CF®488A, CF®633, CF®555 (Biotium, Fremont, CA). Nucleii were stained using 4′, 6-diamidino −2-phenylindole (DAPI) (1:2000, Sigma).

### F-actin sedimentation assay

MGE cell pellets from ten E13.5 embryos, were resuspended in 2 ml of lysis buffer (50 mM PIPES, pH 6.9, 50 mM NaCl, 5 mM MgCl2, 5 mM EGTA, 5% glycerol, 0.1% NP40, 0.1% Triton, 0.1% Tween20, 0.1%-mercaptoethanol), sheared with 25G needle and incubated at 37°C for 10 minutes. Nuclei were removed by centrifugation (2,000 *g*, 5 minutes) and the supernatant subjected to a high-speed centrifugation step (100,000 *g*, 1 hour, 22°C). Pellets (F-actin) were subsequently dissolved in water containing 10 μM cytochalasin D. Equal amounts of each fraction were analyzed by western blot.

### Western Blot

Lysates were run on a 12% SDS-PAGE and transferred into Nitrocellulose membranes (Whatman GmbH, Dassel, Germany). Membranes were subsequently blocked with 5% milk in PBS, 0,1% Tween-20 and immunoblotted with mouse monoclonal anti-actin (Sigma-Aldrich), diluted in blocking solution. Secondary antibodies used: anti-mouse-IgG-Horseradish Peroxidase (GE Healthcare, Buckinghamshire, UK, 1:4000).

### Ion torrent library preparation sequencing

For RNA-seq of Rac1/Rac3wt and Rac1/Rac3dmut samples (3 samples/genotype), total RNA from a pool of MGEs dissected from three E13.5 embryos per sample, was extracted and purified on columns (Qiagen RNeasy Plus Micro kit) and consequently was subjected to mRNA isolation, followed by RNA fragmentation, adaptor hybridization and ligation, reverse transcription, PCR amplification, template preparation and Ion Torrent semiconductor sequencing (3 samples/PI chip, 30-40 M reads/sample). Three biological replicates were used for each condition. Briefly, mRNA isolation was performed with Dynabeads mRNA DIRECT Micro Kit (Life Technologies) according to the manufacturer’s instructions. Library construction was performed with the Ion Total RNA-seq kit (Life technologies) and the Ion XpressRNA kit (Life technologies) was used for barcoding. Amplified libraries were quantified with Qubit (Life Technologies) and Agilent 2100 Bioanalyzer (High Sensitivity DNA kit, Agilent Technologies). Emulsion PCR and enrichment steps were carried out using the Ion PI Hi-Q OT2 200 kit. Sequencing was undertaken using PI chips in an Ion Torrent Proton platform. The Ion PI Hi-Q Sequencing 200 kit (Life Technologies) was used for all sequencing reactions and Torrent Suite™ Software 5.0 (Life Technologies) was used for QC metrics and fastq generation.

### Real time PCR

Extraction of total RNA was performed on samples containing dissected MGEs from E13.5 Rac1/Rac3wt and Rac1/Rac3dmut embryos, each sample consisting of MGEs from 3 embryos, using the RNAeasy Plus Micro kit (Qiagen) according to the manufacturer’s instructions. Total RNA was subjected to reverse transcription according to the protocol of Prime Script 1st Strand cDNA Synthesis kit (Takara Bio USA). Real time PCR analysis was performed using a Step One Plus real-time PCR system (Applied Biosystems, Life Technologies, Thermo Fisher Scientific Inc., Waltham, MA). Expression levels of genes encoding Ank-1 (forward primer (F): 5’-ATCAAAGACATCGAGGCGCA-3’, reverse primer (R): 5’-ACCCGTTCTGGTTACAGGTG-3’), Ccnl1 (F: 5’-GGAAAAAGGACTCCAAGCCC-3’, R: 5’-CCCAACTCCTTTAGCACCCT -3’), Dyntl1 (F: 5’-GGCGAAGGAGATGCGTTAAA -3’, R: 5’-TGGCGCTTTCTATAGCCTCC -3’), Hist2h2ac (F: 5’-ACCTCGGTGTGTCCTACATTA-3’, R: 5’-GCGTAGTTGCCCTTGCG-3’), Magel2 (F: 5’-AAGGGCACTCAACCATTCGT-3’, R: 5’-GACCTGTCGGATCAGTGGTG-3’), Meg3 (F: 5’-GGGTAGGCTCAGCATGGTTT-3’, R: 5’-CCCGCTCCAGATTTCACCTT-3’), miR-124 (F: 5’-GAAGGATCGACCCAGCACAA-3’, R: 5’-TAAGGGCTACTTTGCGAGGC-3’), Miat (F: 5’-GGGCTTAGGGGAGTCCAAAC-3’, R: 5’-CTCACTACCAACCCCAACCC-3’), Sptb (F: 5’-GAACATTGACGCAGAGGGGA-3’, R: 5’-AGTTGTTCTGCCTTCGCCTT-3’), Tpcn2 (F: 5’-GGCCTGTTACATTGGTGGGA-3’, R: 5’-TGCAGCACCGGAGTGGAC-3’), Zmym6 (F: 5’-TTCTCTCGGCTTTCAGGTTGG -3’, R: 5’-TCCGGTAGGTAGGTTGTCCC -3’) were examined. As an internal control GAPDH was used (F: 5’-ATTGTCAGCAATGCATCCTG-3’, R: 5’-ATGGAC TGTGGTCATGAGCC-3’). Real-Time PCR was performed in a 15 μl reaction containing 7.5 μl of KAPA SYBR FAST qPCR Master Mix, 1.5μl forward primer (2 pmol/μl), 1.5μl reverse primer (2 pmol/μl), 3.5μl cDNA sample, 0.3μl ROX High Reference Dye (50X) and 0.7μl water according to the manufacturer’s instructions. PCR conditions were 95°C for 20sec, followed by 40 cycles of 95 °C for 3sec, 58 °C (Ccnl1, Dyntl1, Hist2h2ac, Magel2, Meg3, MiR124, Miat, TPCN2, Zmym6) /57 °C (Ank1, Sptb) for 30sec and 72 °C for 1sec, and a final step of 95 °C for 15sec, 60°C for 1min and 95°C for 15sec. We performed comparative real-time PCR analysis for Ank1, Ccnl1, Hist2h2ac, Magel2, Meg3, miR-124, Miat, Sptb, and a relative standard curve was performed for Dntl1, TPCN2, Zmym6. PCR runs were performed for each sample in triplicates using cDNA corresponding to 1ng of total RNA regarding Ccnl1, Hist2h2ac, 5ng regarding Dyntl1, Magel2, Meg3, MiR124, Miat, TPCN2, 10ng regarding Zmym6, and 50 ng Ank1, Sptb. Expression levels for all genes were normalized according to the internal control and expressed as percentage of control change.

### Time lapse imaging

Imaging was performed on LeicaSP8 confocal microscope. All live cells were imaged on Leica SP8 confocal microscope enclosed in an environmental chamber with temperature and CO2 control. Images of living YFP expressing MGE-derived interneurons migrating in E14.5 Rac1/Rac3wt and Rac1/Rac3dmut embryonic brain slices or RFP expressing ones, upon focal electroporation were acquired every 3min, for up to 8h at the level of the corticostriatal boundary or in the SVZ of the MGE respectively. Images of living YFP expressing, Rac1/Rac3 wt interneurons treated with trans Ned-19 and DMSO were acquired for up to 8h every 6min. Images of MGE-derived interneurons plated in matrigel were collected every 3 min for up to 8h using a transmitted light detector. Images of living Rac1/Rac3wt and Rac1/Rac3dmut interneurons migrating from MGE explants stained with the fluorescent dyes lysotracker Red DND and Hoechst were acquired every 3 min for 4h. In all the cases above successive “z” optical plans were collected with a step size of 1μm. Lastly, images of YFP-expressing interneurons, Rac1/Rac3wt and Rac1/Rac3dmut stained with lysotracker Red DND and tubulin tracker deep red were captured every 523ms for 60sec with resonant scanning xyt acquisition mode.

### Data analysis

Gene Ontology (GO) was performed using Cytoscape software (Otasek et al., 2019; Shannon et al., 2003). All image analysis was performed with ImageJ, in its standard version (http://rsbweb.nih.gov/ij/; RRID: nif-0000-30467) or in the Fiji distribution (http://fiji.sc/Fiji; RRID: SciRes_000137) (Schindelin et al., 2009; Schneider et al., 2012). The time-lapse data were assembled and analyzed by Fiji. Parameters such as velocity, duration of the migratory cycle, amplitude and frequency of nuclear translocation were analyzed using Mtrack J plugin (Meijering et al., 2012). Intensity of fluorescent signal in standard distances from the center of the nucleus to the tip of the principal neurite was quantified with the Concentric Circles plugin. The radii of the concentric circles were: 1^st^ circle radius: 1.20 μm, 2^nd^: 2.15, 3^rd^: 3.10, 4^th^: 4.05, 5^th^: 5.00, 6^th^: 5.96, 7^th^: 6.91, 8^th^:7.86, 9^th^:8.81, 10^th^:9.77, 11^th^:10.72, 12^th^: 11.67, 13^th^:12.62, 14^th^:13.57, 15^th^: 14.53, 16^th^: 15.48, 17^th^: 16.43, 18^th^: 17.38 μm, 19^th^: 18.34, 20^th^: 19.29 and 21^th^: 20.24 μm. Fluorescence intensity time plot of migrating interneurons stained with Hoecht and Lysotracker red were created using the ImageJ macro-script, Intensity2 (author Steve Rothery, FILM-Facility for Imaging in Light Microscopy, Imperial College London https://www.imperial.ac.uk/medicine/facility-for-imaging-by-light-microscopy/software/fiji/). Analysis of TPC2 expression was performed using the 3D object counter plugin measuring fluorescent puncta per cell in thresholded images (Bolte and Cordelières, 2006). Kymograph were created with the Multi Kymograph plugin on Fiji (authors J.Reitdorf, A.Seitz) and were analyzed using the semi-automated software package Kymoanalyzer v101 (Neumann et al., 2017). The morphology of the interneurons was analyzed with the semi-automated plugin SNT-Simple Neurite Tracer (Arshadi et al., 2021). The Sholl analysis was performed using the Sholl plugin of Fiji (Ferreira et al., 2014). Principal neurite length in DMSO or trans Ned-19 treated interneurons was quantified using the plugin AnalyzeSkeleton (Arganda-Carreras et al., 2010).

### Statistical analysis

All data are presented as mean ± SEM. Statistical analysis of the measured values for each group was performed using two-tailed, unpaired Student’s t-test or One-way analysis of variance ANOVA followed by Tukey’s or Sidak’s post-hoc tests for multiple comparisons. Also, multiple tests were used followed by two-stage linear step-up procedure of Benjamini, Krieger and Yekutieli. Statistical analysis was performed using GraphPad Prism version 8.00 for Windows (Prism, GraphPad). P-values less than 0.05 were considered to be statistically significant.

## ACKNOWLEDGEMENTS

We would like to sincerely thank the animal facility and the Genomics Facility, IMBB FORTH, Heraklion, Crete, Greece. We are grateful to Dr Ivan de Curtis for providing the Rac3 mutant line, Martha Gjikolaj for help with slice electroporation and cultures and Dr Marina Vidaki for constructive comments on the manuscript.

## FUNDING SUPPORT

We are grateful to the Manasaki Foundation for providing scholarship funds for ZK through the UoC, the progrma EΔΒΜ34-new researchers supported by the Operational Program “Human Resources Development, Education and Lifelong Learning” which is co-financed by the European Union (European Social Fund) and Greek national funds and the Hellenic Foundation for Research and Innovation (Grant 1621) to the Karagogeos lab. DK was supported by the Luxembourg National Research Fund (FNR) through PRIDE17/12244779/PARK-QC. Financial support to MD was provided by the Hellenic Foundation for Research and Innovation (Grant 2564), Fondation Santé, and a Stavros Niarchos Foundation grant to B.S.R.C. “Alexander Fleming”, as part of the Foundation’s initiative to support the Greek research center ecosystem.

The authors declare no conflicting interests.

The RNAseq data can be found at:

https://www.ncbi.nlm.nih.gov/geo/query/acc.cgi?acc=GSE206752

