## Supplemental Figures for "Rac1, Rac3 GTPases and TPC2 are required for axonal outgrowth and migration of cortical interneurons"

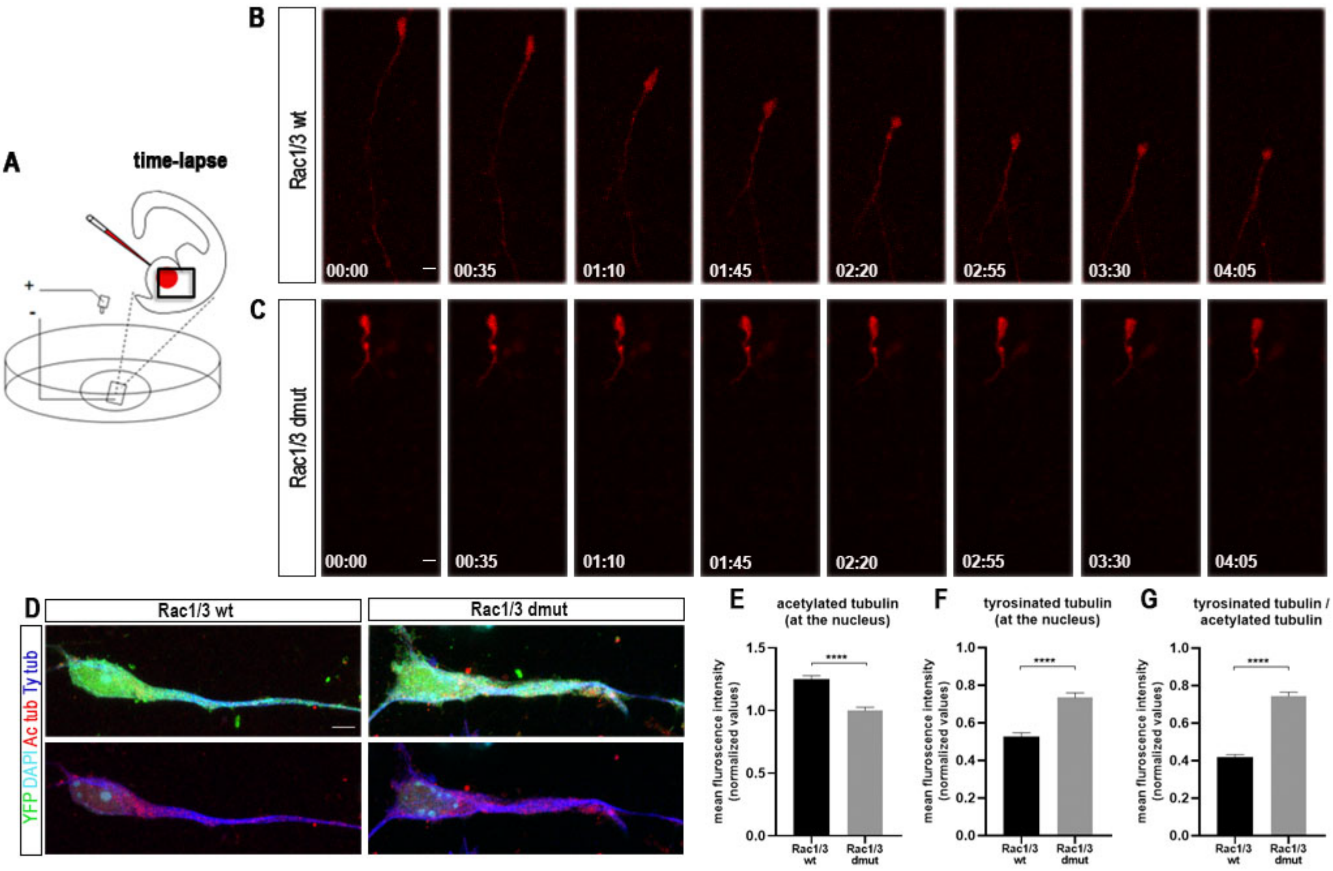


**Sup fig. 1 Impairments in nucleokinesis in MGE-derived interneurons due to the absence of Rac1/ Rac3.**

(A-C) Impaired motility of interneurons in the region of MGE. (A) schematic representation of the focal electroporation of a plasmid solution (red) in the MGE of E14.5 cultured brain slice. Boxed area indicates the imaging window. Time-lapse sequence of an electroporated Rac1/3wt interneuron (B) and a Rac1/3dmut interneuron (C) (D-F) Impairments in microtubule dynamics. (D) Immunostaining for ac-tubulin (red), ty-tubulin (blue), YFP (green) in the perinuclear region of interneurons exiting from MGE explants, 2DIV. Graph illustrating intensity values (E) of ac-tubulin (F) ty-tubulin and (G) ty-tubulin/ac-tubulin ratio. (n=40-45 cells per condition, from 3 embryos). Student's t test****P < 0.0001. Error bars indicate SEM. Scale bars represent 4 μm.

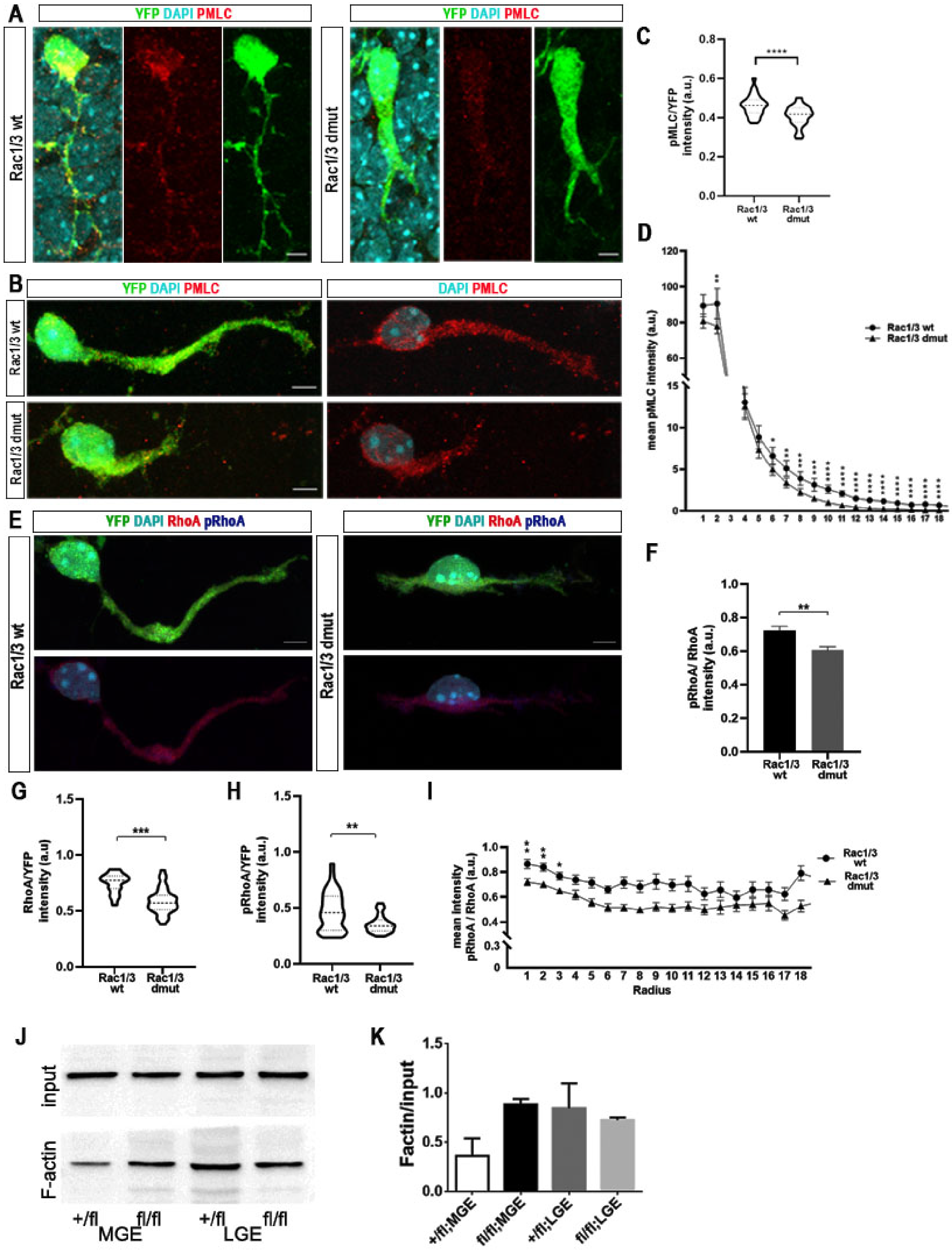


**Sup fig 2 Impairments in actomyosin dynamics in interneurons upon ablation of Rac1/Rac3.**

(A,B) Immunostaining showing the subcellular distribution of phospho-Myosin light chain (PMLC, in red) and YFP (green) in migrating Rac1/3wt and

Rac1/3dmut interneurons, at E14.5 brain slices (A) and in cultured MGE-derived interneurons, 3DIV (B). The nucleus is labeled with DAPI (cyan). (C) Graph illustrating normalized intensity values of pMLC (F) and (D) mean intensity values of PMLC measured in standard radii concentric circles starting from the center of the nucleus. (E) Immunostaining for RhoA (red), pRhoA(blue) and YFP (green) in cultured MGE-derived interneurons from E14.5 Rac1/3wt and Rac1/3dmut embryos. The nucleus is labeled with DAPI (cyan). (F-I)) Quantification analysis of the intensity of the proteins. Graph illustrating (F) pRhoA/RhoA ratio of the intensity values, (G) normalized intensity values of RhoA (H) pRhoA and (I) pRhoA/RhoA measured in standard radii concentric circles, starting from the center of the nucleus. (n=30 cells per condition, from 3 embryos). (J-K) F-actin fraction is increased in Rac1-mutant. Quantification of the F-actin isoform by sedimentation assay, using anti-actin western blot on protein extracts from E13.5 MGE, indicated the same amount of total actin (input) and different F-actin. Student's t test (C, F, G, H), multiple t test (D,I), $*P<0.05$, $**P<0.01$ $***P<0.001$, $****P < 0.0001$. Error bars indicate SEM. Scale bars represent 4 μm.

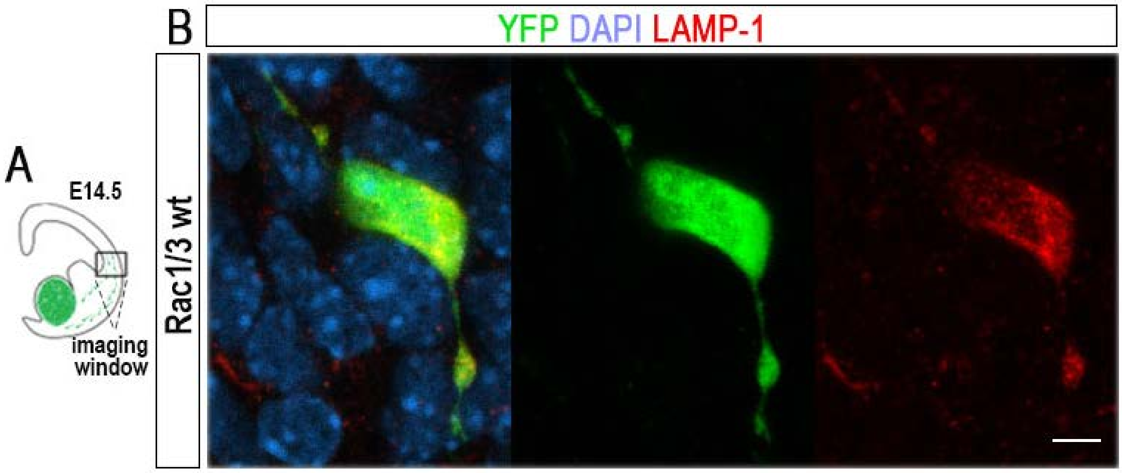


**Sup fig. 3 Lysosome distribution in migrating interneurons.**

(A) Schematic representation of E14.5 brain slice. (B) Immunostaining for late lysosomes (LAMP-1, in red), YFP (green) in migrating Rac1/3wt interneurons, at E14.5 brain slices. Scale bars represent 4 μm.

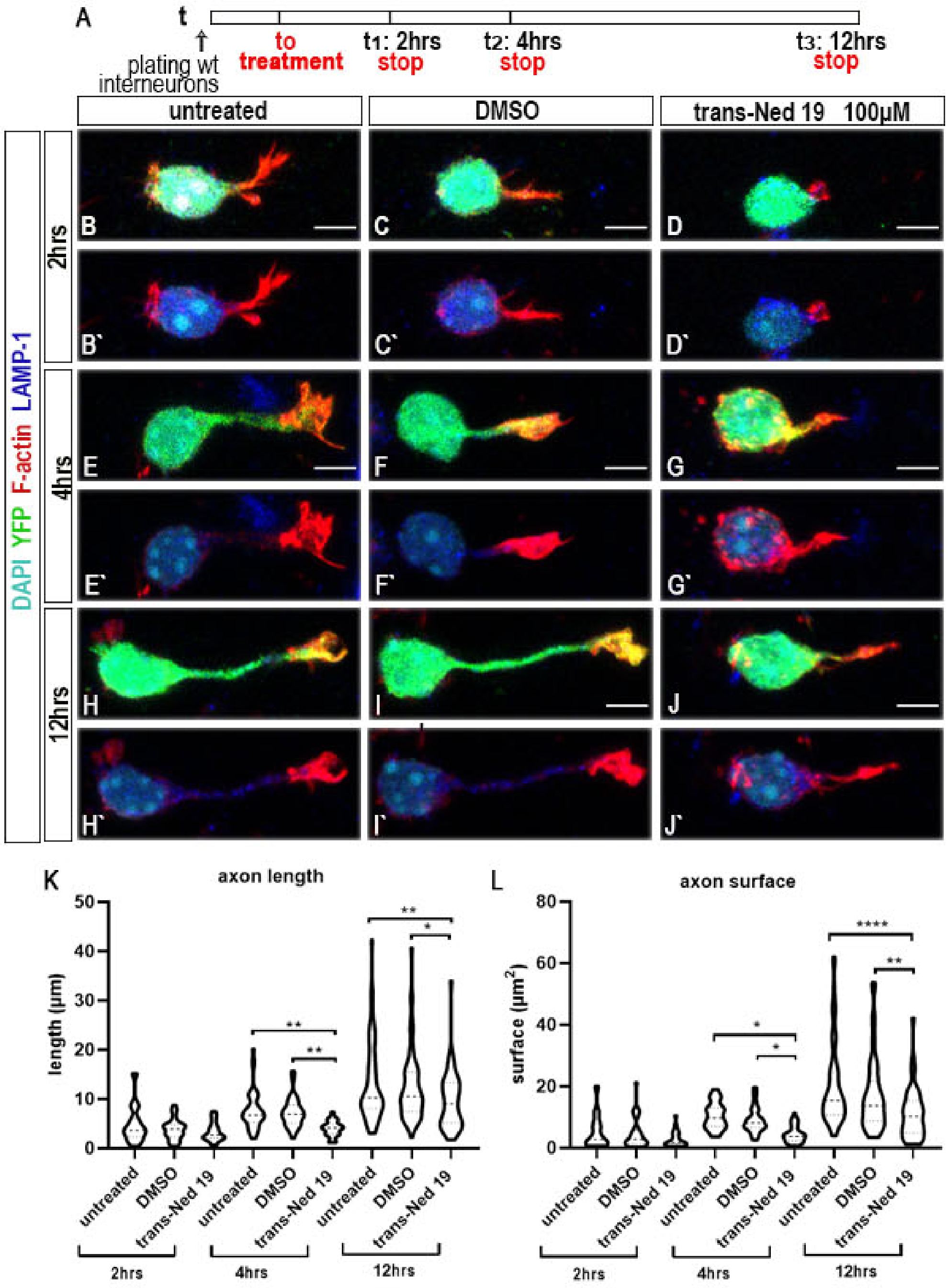


**Sup. Fig 4 Pharmacological inhibition of TPC2 negatively affects axon outgrowth initiation of wild type interneurons.**

(A) Schematic representation of the treatment protocol. (B-J') Immunostaining for F-actin (phalloidin, in red), YFP (green), late lysosomes (LAMP-1 in blue), in cultured MGE-derived interneurons from E14.5 Rac1/3wt embryos. The nucleus is labeled with DAPI (cyan). Interneurons were untreated (B-B', E-E', H-H') or were treated with DMSO (C-C', F-F', I-I') and trans-Ned 19 (D-D', G-G', J-J') at 100μM concentration for 2hrs (B-D'), 4hrs (E-G') and 12hrs (H-J'). Graphs indicating principle neurite length (K) and principle neurite surface (L) (n=35-45 cells per condition, from 3 independent experiments). One-way ANOVA, Tukey's multiple comparisons test, *$P<0.05$, **$P<0.01$, ****$P < 0.0001$. Error bars indicate SEM. Scale bars represent 4 μm.

**Movie 1**: Migration of MGE-derived interneurons.

**Movie 2**: Lysosomal positioning in migrating interneurons.
